# *Streptomyces* enrichment in roots during drought is uncoupled from plant benefit and is driven by host suppression of iron uptake and immunity

**DOI:** 10.64898/2026.03.06.710171

**Authors:** Connor R Fitzpatrick, Ryker Allen Smith, Junko Hige, Theresa F Law, Dor Russ, Oluwadamilola Elizabeth Ajayi, Abdul Aziz Eida, Pierre Jacob, Monet Jowers, Narender Kumar, Cindy Thao Uyen Lai, Manuel Anguita-Maeso, S. Brook Peterson, Chinmay Saha, Tara Skelly, Qinqin Zhao, Wenbin Zhou, Sarah R Grant, Joseph D Mougous, Corbin D Jones, Jeffery L Dangl

## Abstract

Drought reshapes plant root microbiota, yet the mechanistic drivers and consequences of this observation remain unclear. We discovered that suppression of host immunity and iron homeostasis is required for *Streptomyces* enrichment in roots during drought across diverse soils. Genetic and physiological manipulation of these host pathways confirmed their requirement in modulating *Streptomyces* root enrichment. Drought-induced suppression of iron uptake was conserved across the ∼160 My monocot-eudicot divergence. Some *Streptomyces* strains enhanced plant growth and rescued iron uptake under drought. These benefits were uncoupled from *Streptomyces* root enrichment. They were instead shaped by intra-*Streptomyces* antagonism. We propose a two-step model: drought-driven down regulation of host iron and immune pathways enriches *Streptomyces*, while intra-genus dynamics fine-tune strain-level assembly and functional outcomes. Our data refine the idea that *Streptomyces* are enriched in roots during drought in response to a plant ‘cry for help’ and consequently contribute to alleviation of this abiotic stress.

## Introduction

Plants assemble distinct microbial communities (microbiota) from their environments that vary in diversity and composition across plant species, organs, epi- and endophytic habitats, and soil locality^1–7^. The primary determinant of plant microbiome composition is the resident soil microbial community, followed by plant organ identity, such as leaf or root^8,9^. These observations led to a multistep model of community assembly in which plant habitats, from epi- to endophytic compartments, sequentially winnow the diverse soil microbiota into a selected community^9^. This process reflects both deterministic and stochastic ecological factors, including plant physiological and immune conditions and extensive microbe–microbe interactions within plant-associated habitats^10–13^.

The functional importance of various plant microbiota is indisputable. Stereotypic plant development, immune function, and nutritional programs are deeply integrated with associated microorganisms^13–20^. Perturbation of the ‘typical’ microbiome assembly can lead to dysbiosis^19–21^ and environmental change can reshape host plant microbiome composition^22^. These observations suggest that plant-associated microbiota are tightly regulated by the plant to maintain host health^19–21^. This concept led to the ‘cry for help’ hypothesis wherein plants actively shape their microbiome in response to environmental perturbations to benefit from the enrichment or depletion of particular microorganisms^22–28^.

During drought, members of the Actinomycetota, in particular *Streptomyces*, become enriched in plant roots, but not in surrounding soil^29–33^. *Streptomyces*, a ubiquitous group of soil bacteria and prominent member of the plant microbiome, arose 450 Mya, following the migration of plants to land^34^, and are well-studied for their secondary metabolite production^35,36^. The drought enrichment of *Streptomyces* is conserved across host plant species spanning 350 My of divergence^29,32,37,38^, suggesting a conserved drought response in plant roots that favors *Streptomyces* enrichment. The near ubiquitous enrichment of *Streptomyces* under drought has provoked the hypothesis that they are responding to a ‘cry for help’ from the plant and are involved in alleviating host drought stress^31–33,39,40^.

Despite years of investigation, the mechanistic drivers and functional consequences of *Streptomyces* enrichment in roots under drought remain poorly understood ^39,41^. We show here that rather than positively selecting *Streptomyces*, drought suppresses host plant immunity and iron homeostasis that otherwise inhibit the proliferation of *Streptomyces* in plant roots. We also show that certain *Streptomyces* strains promoted plant growth and maintained iron uptake during drought. However, these benefits were not linked to enrichment; instead enrichment among *Streptomyces* was governed by competition among strains. Thus, under drought, iron-stressed and immune-suppressed roots become a niche exploited by *Streptomyces*. Intra-genus competitive success is uncoupled from plant-beneficial traits leading to inconsistent effects on plant health, incompatible with the simple ‘cry for help’ hypothesis.

### *Streptomyces* are enriched in roots during drought across diverse soils

We imposed chronic drought stress on *Arabidopsis thaliana* (Col-0) grown in soils collected from 18 locations across the USA (Figure 1A). These sites span the climatic range of the region (Figure S1A), and the soils vary widely in physical and chemical properties (Figure 1B, Table S1). After 2 weeks under benign conditions, plants were maintained for 6 weeks under standardized drought or watered regimes corresponding to gravimetric water contents at permanent wilting point or field capacity, respectively, determined by water retention analysis (Figure S1B). On average, drought reduced biomass 4.3-fold (range across soils: 2–10.5-fold), whereas biomass under watered conditions varied 20-fold across soils (Figure S1C).

**Figure 1.**
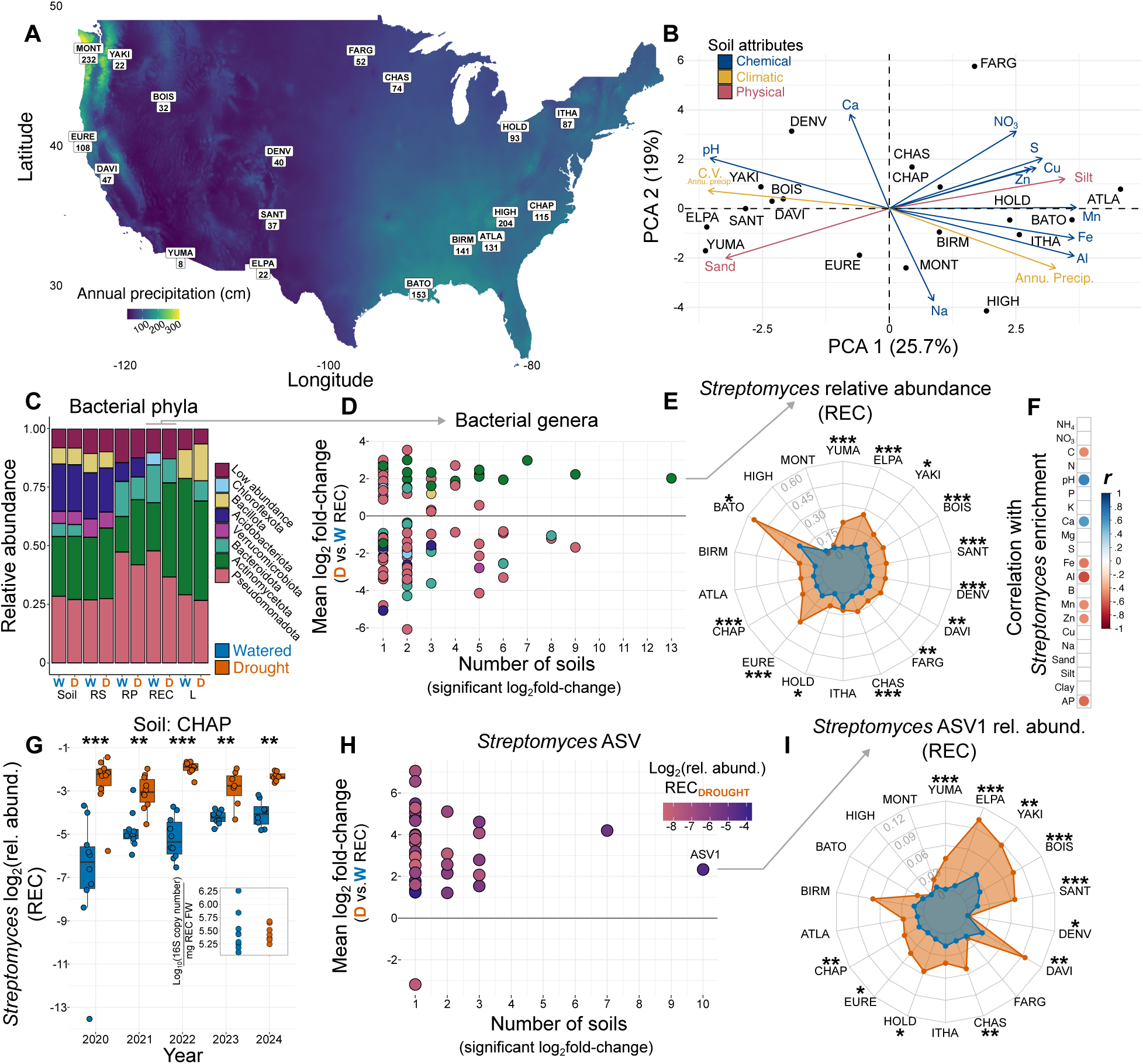
Streptomyces are uniquely enriched in plant roots during drought across diverse soils. (**A**) Soil locations denoted with a four-letter code and annual precipitation. (**B**) A PCA biplot showing the dispersion of soil chemical and physical attributes and the climatic attributes (vectors) of each location. (**C**) Relative abundance of bacterial phyla averaged across all soil locations in different habitats under watered and drought conditions (rhizosphere, RS; rhizoplane, RP; root endophytic compartment, REC; leaf surface and endophytic compartment, L; *n* = 7-20 per location X habitat X treatment combinations). (**D**) Bacterial genera vary in their differential abundance (log_2_ fold-change, LFC) in response to drought in the REC. The number of soils in which significant differential abundance was observed for a given genus is shown along the x-axis (*n* = 7-20). (**E**) The relative abundance of the most prevalently enriched bacterial genus, *Streptomyces,* in the REC under watered and drought conditions in each soil location. Note that locations are ordered clockwise from dryest (YUMA) to wettest (MONT). (**F**) Pearson correlation coefficients between the *Streptomyces* REC LFC and soil and climate attributes across locations (*n* = 18). (**G**) Independent experiments using soil sourced from the same location (CHAP) demonstrate that *Streptomyces* enrichment in roots during drought is temporally consistent (*n* = 8-12). (**G** inset) The bacterial load, estimated from 16S copy number, does not decrease in the REC during drought (*n* = 10, two-tailed t-test). (**H**) *Streptomyces* ASVs vary in their differential abundance (log_2_ fold-change, LFC) in response to drought in the REC. The number of soils in which significant differential abundance was observed for a given ASV is shown along the x-axis. Point colors correspond to the average relative abundance of individual *Streptomyces* ASVs. (**I**) The relative abundance of the widespread and highly abundance *Streptomyces* ASV1 in the REC under watered and drought conditions in each soil location (*n* = 7-20). Significance in EGI determined by Wald tests with FDR adjusted p values. For G (and all subsequent boxplots presented) the horizontal bars within the boxes represent the medians. The tops and bottoms of the boxes represent the 75th and 25th percentiles, respectively. The upper and lower whiskers represent 1.5× the interquartile range from the upper edge and lower edge of the box. Statistical significances in all figures indicated by ∗∗∗ p < 0.001, ∗∗ p < 0.01, ∗ p < 0.05.

We sampled bacterial microbiota from bulk soil, leaves (combined epi- and endophytic habitats), and roots, which were further separated into rhizosphere (RS), rhizoplane (RP), and root endophytic compartment (REC). The V3–V4 region of the bacterial 16S rRNA gene was sequenced and reads were assigned to sequence-identical amplicon sequence variants (ASVs) for community analysis. Resident soil bacterial communities were highly diverse with little overlap in individual ASVs across locations (Figure S1D), resulting in strong effects of soil locality on plant microbiome diversity and composition, followed by plant habitat fraction (Figure S1D and S1F). In contrast, drought produced no consistent shifts in microbiome composition or diversity across soils (Figure S1D and S1F), instead generating soil-specific responses (Figure S1E and S1G). Despite this variation, Actinomycetota were consistently enriched in root-associated communities, specifically the RP and REC (Figure 1C), across all locations. This enrichment was absent in soil and leaf communities (Figure 1C), indicating that drought-driven shifts are confined to habitats closely associated with roots.

We analyzed REC community composition at higher taxonomic resolution and found that the genera most consistently enriched belonged to Actinomycetota, with *Streptomyces* showing the strongest and most widespread enrichment (Figure 1D). However, enrichment varied across soils (Figure 1E) and correlated with environmental attributes including annual precipitation and soil pH (Figure 1F). We next tested whether enrichment was temporally consistent across soils. *Streptomyces* log₂ fold-change values were positively correlated among soil locations across two independent experiments (*r* = 0.66, *p* = 0.003), indicating reproducible enrichment. Similarly, independent experiments using soils collected from a single site across five consecutive years produced comparable *Streptomyces* enrichment in the REC during drought (Figure 1G).

Enrichment could reflect either increased growth or persistence during a broader decline of the bacterial community. To distinguish these possibilities, we used estimated 16S copy number^42^ as a proxy for bacterial abundance and detected no change in REC copy number during drought, indicating that persistence alone does not explain enrichment (Figure 1G inset). Together, these results show that *Streptomyces* enrichment is geographically widespread, temporally stable, and driven, at least in part, by growth in plant roots during drought.

Individual *Streptomyces* ASVs varied widely in both the magnitude and prevalence of enrichment (Figure 1H). Most ASVs were enriched at only one or a few locations (Figure 1H), reflecting their limited distribution across soils. In contrast, ASV1, a widespread plant-associated *Streptomyces* clade, was enriched at 10 of the 17 locations where it occurred (Figure 1I). The prevalence of *Streptomyces*—particularly ASV1—enrichment is notable given the large variation in bacterial diversity and community composition across soils. However, enrichment was not universal, despite being the most frequent abundance shift in the root microbiome during drought. These patterns suggest that *Streptomyces* enrichment is shaped by interactions among soil properties, resident microbial communities (including local *Streptomyces* populations), and plant responses to drought. Our results further indicate that shared mechanisms likely promote enrichment across soils, although their effects vary among locations.

### Suppression of plant immunity and iron uptake during drought accompanies *Streptomyces* enrichment in plant roots

Despite being one of the most widely observed stress-induced shifts in host-microbiota composition, the mechanisms underlying *Streptomyces* enrichment in plant roots during drought and how they relate to plant health remain poorly understood^39^. Using transcriptional profiles of roots and shoots, we identified specific plant responses to drought across different soils that associate with variable *Streptomyces* enrichment. Although many drought responsive genes (DRGs) were observed in individual samples, fewer genes were commonly differentially expressed in roots or shoots of plants grown in multiple soil samples (Figure 2A, Table S3). We binned these DRGs into cloud, shell, and core, representing DRGs that were found in few (1-6), intermediate (7-13), and most soils (14-15), respectively. We observed heterogeneity in gene expression across soils, highlighting the complexity of plant drought responses in natural soils, yet a readily observable core transcriptional response to drought occurs (Figure 2B and S2A). As expected, biological processes related to water stress were enriched in core and shell upregulated DRGs in roots (Figure 2B), including genes responsive to abscisic acid (ABA), the chief plant hormone activated during drought^43^ (Figure S2B). We tested whether such prevalent responses to drought were contributing to *Streptomyces* enrichment by comparing wild type Arabidopsis with mutants perturbed in the biosynthesis and response to ABA in CHAP (formerly, MF or Mason Farm) soil. Interestingly, we failed to find any attenuation of the *Streptomyces* enrichment in these mutants (Figure S2C), indicating that prevalent drought responses governed by ABA are not responsible for the widely observed REC enrichment of *Streptomyces* during drought.

**Figure 2.**
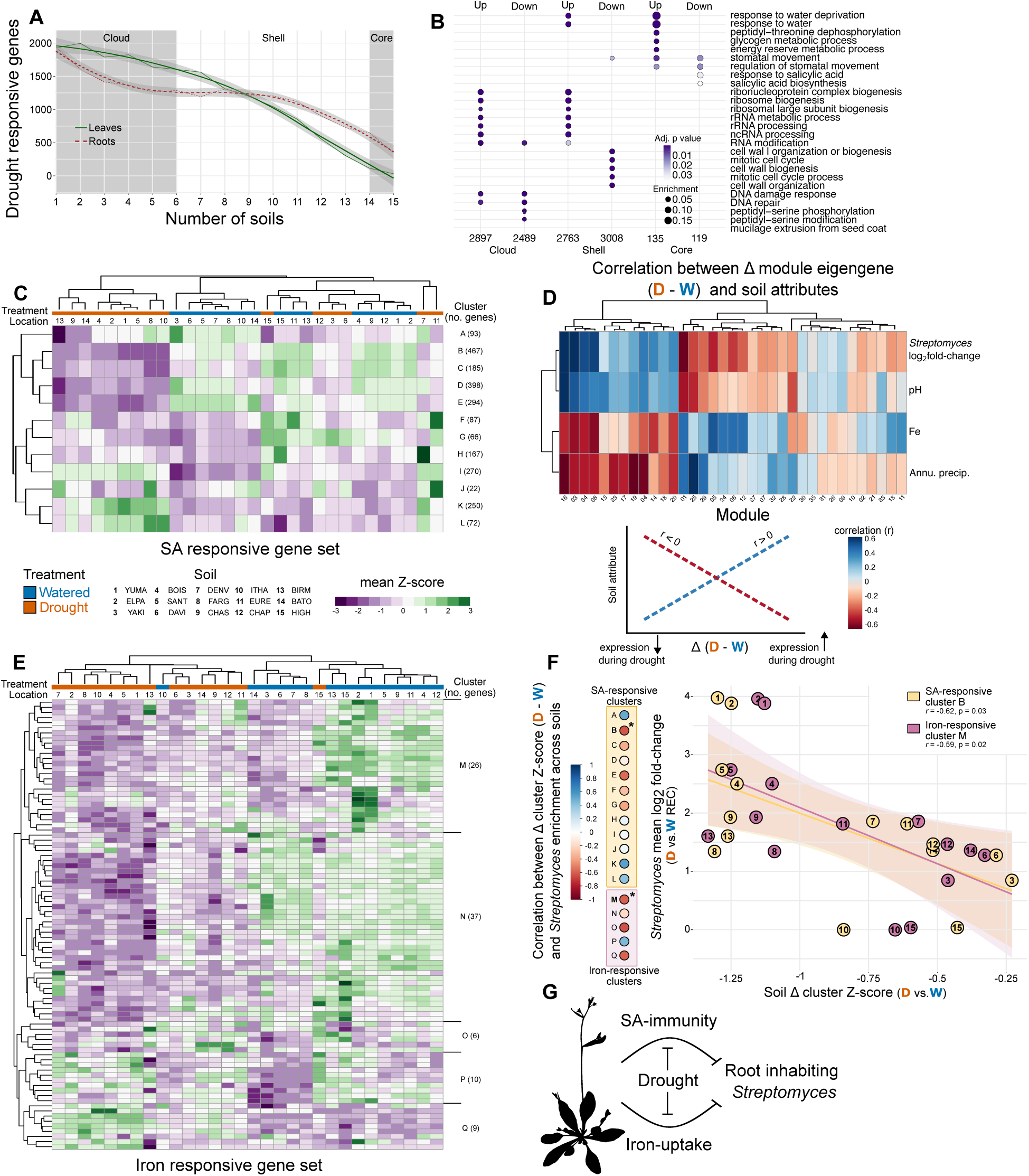
Drought suppresses subsets of SA and iron responsive genes. (**A**) Drought responsive genes (fold change > 1.5, adjusted P value < 0.05) in leaves and roots across the 15 soil locations selected for transcriptional profiling (*n* = 3-6 per location X organ X treatment combination). (**B**) Enrichment analysis of biological processes in up- and downregulated genes in the cloud, shell, and core drought responsive genes. (**C**) The expression in plant roots (average z-score) of a literature derived set of SA responsive genes (defined in leaves) under watered and drought conditions across all soils. Each row represents a gene cluster defined from a hierarchical clustering analysis of the expression of individual genes in the SA responsive gene set in our samples. (**D**) Weighted gene co-expression network analysis (WGCNA) modules of genes with similar co-expression profiles (heatmap columns). We correlated location-specific differences in the expression (eigengenes) of these modules in roots under drought vs. watered conditions with soil attributes (heatmap rows), including the LFC of *Streptomyces* in roots during drought (see bottom of panel D for a graphical representation). (**E**) The plant root expression (average z-score) of a literature derived set of iron responsive genes (defined in roots) under watered and drought conditions across all soils. Each row represents an individual gene in the iron response gene set. (**F**) The differential expression of subsets of the SA- (clusters A-L) and iron-responsive gene sets (clusters M-Q) are correlated with *Streptomyces* enrichment in roots during drought (*n* = 15). (**G**) During watered conditions, plant iron uptake and SA-related immunity inhibit *Streptomyces.* Under drought, plant iron uptake and SA-related immunity are suppressed, which contributes to the enrichment of *Streptomyces*.

Downregulated DRGs, in the cloud and shell, were enriched for biological processes related to peptidyl-serine modifications, DNA repair, and cell wall biogenesis (Figure 2B), suggesting suppression of these functions occurs in a soil specific manner. Consistent with prior work^44–46^, salicylic acid (SA) biosynthesis and responses were among the few biological processes enriched in the downregulated core DRGs (Figure 2B). SA is a plant hormone that orchestrates many facets of plant immunity in both local and systemic tissues^47–49^ and modulates the assembly of the root microbiome^50^. Using a literature-derived set of SA responsive genes (Table S4), we discovered several gene clusters that exhibited suppression in roots during drought, while other SA-responsive gene clusters had the opposite regulation (Figure 2C). Given the role of SA-dependent responses in plant-microbe interactions, we hypothesized that its regulation during drought could contribute to *Streptomyces* enrichment.

We performed weighted gene co-expression analysis^51^ to define modules of genes based on similar co-expression patterns to directly associate the drought enrichment of *Streptomyces* in roots with both root transcriptional responses and soil properties. We correlated the difference in the average expression of these gene modules (eigengene) between drought and watered conditions across soils with observed values of *Streptomyces* REC enrichment and soil attributes (Figure S2D and S2E). We observed a striking pattern: gene modules whose regulation was strongly associated with *Streptomyces* enrichment were similarly positively associated with soil pH and inversely associated with iron availability (Figure 2D, S2D, and S2E). This suggested that *Streptomyces* enrichment is associated with genes whose suppression in roots during drought increases at elevated pH but decreases as soil iron becomes more available. Given the established negative relationship between soil pH and iron availability^52,53^, we analyzed the regulation of iron responsive genes in response to drought across soils (Table S4). Like SA responsive genes, we observed clusters of iron responsive genes that exhibited widespread suppression in roots during drought across soils (Figure 2E).

Plant iron uptake activity and homeostasis were recently linked to root microbial assembly^41,54–57^. Thus, we hypothesized that suppression of host iron-responsive genes in roots during drought contributes to *Streptomyces* enrichment. Indeed, differential regulation of both SA- and iron-responsive genes in response to drought across sites was related to *Streptomyces* enrichment. We found two clusters, SA responsive gene cluster B and iron responsive gene cluster M, whose suppression during drought was strongly correlated with *Streptomyces* enrichment (Figure 2F). Cluster B includes well characterized genes involved in systemic acquired resistance such as *FMO1* and *SARD4*, and cluster M includes critical genes involved in iron acquisition such as *F6’H1*, *BGLU42*, *IRT3*, and *NAS1*. This suggests that SA immunity and iron uptake inhibit *Streptomyces* colonization in plant roots under watered conditions, while during drought the suppression of these processes leads to an increase in *Streptomyces* abundance (Figure 2G).

### Manipulation of plant immunity and soil iron availability modulates *Streptomyces* abundance in roots

Our model (Figure 2G) predicts that manipulation of SA-dependent immunity or iron uptake will change *Streptomyces* abundance in plant roots. To test this model, we first severed the antagonistic link between drought and SA response by foliar application of the SA analog, benzothiadiazole^58,59^ (BTH) to plants grown under watered and drought conditions in CHAP soil (Figure 3A). BTH application resulted in decreased plant biomass (Figure S3A) and elevated expression of the canonical SA response marker *PR1* under watered and drought conditions (Figure S3B). We observed a decrease in *Streptomyces* abundance in roots during both watered and drought conditions following the foliar BTH application (Figure 3B). BTH had little direct effect on *Streptomyces* growth in vitro (Figure S3E), and foliar application resulted in minimal effects on root bacterial community diversity and composition (Figure S3C and S3D). Previous studies reported inconsistent effects of SA on plant-associated *Streptomyces*, reflecting variation among strains and experimental approaches^50,60^. In this work, we chemically modulated endogenous SA responses to avoid the confounding pleiotropic effects associated with constitutive SA-producing mutants^61–64^. Our results are consistent with the endogenous suppression of SA-associated immune responses during drought contributing to the increase in *Streptomyces* abundance in roots.

**Figure 3.**
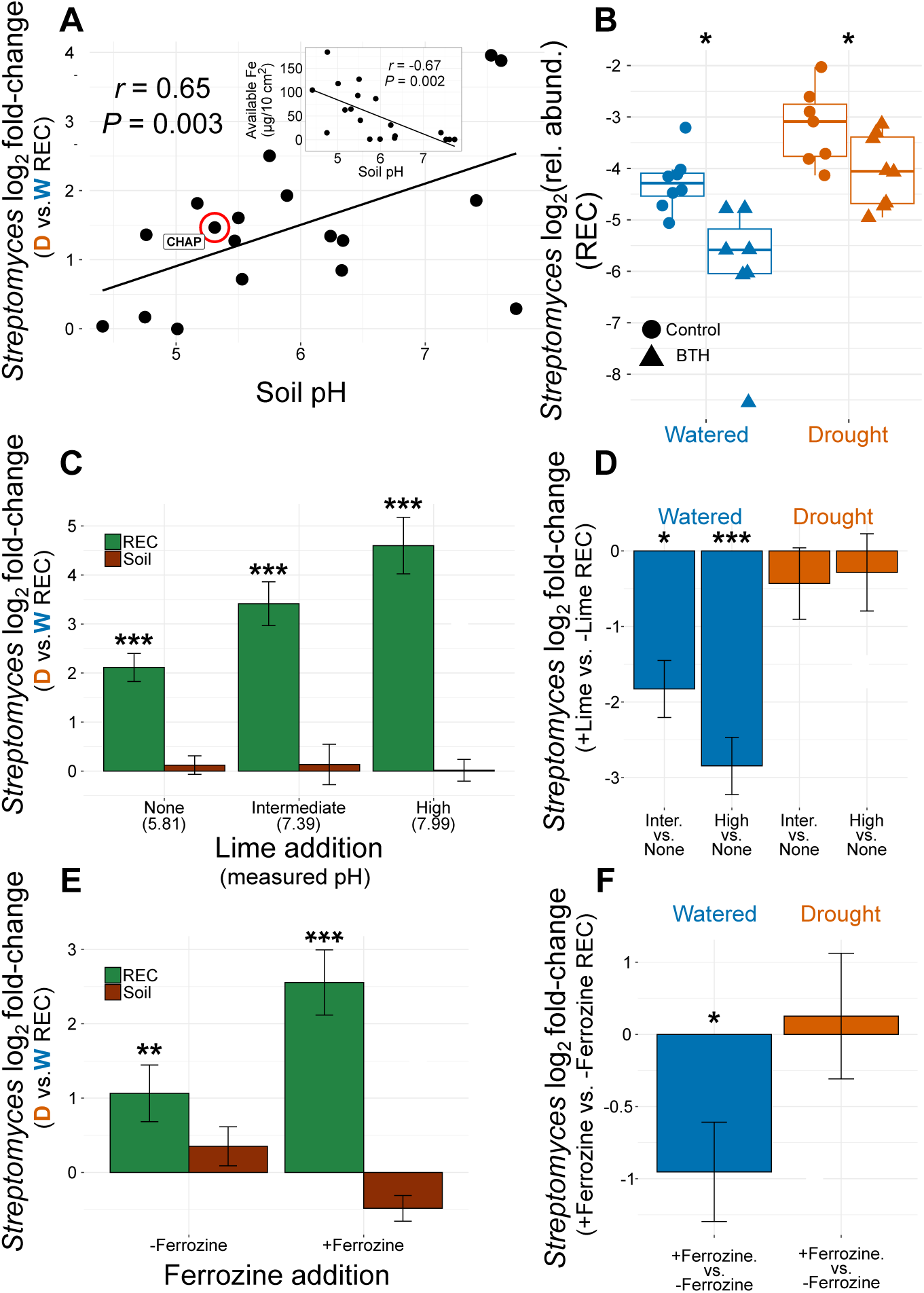
Manipulation of plant immunity and iron-uptake responses modulates *Streptomyces* abundance in roots. (**A**) *Streptomyces* enrichment in roots during drought is positively related to pH across soils (*n* = 18). (**B**) Foliar application of SA mimic, benzothiadiazole (BTH), suppresses *Streptomyces* abundance in plant roots during drought and watered conditions (*n* = 7-8). (**C**) Artificially raising soil pH through the addition of lime results in increased enrichment of *Streptomyces* in roots but not soil during drought (*n* = 8). (**D**) Increased enrichment at elevated pH is caused by a depletion of *Streptomyces* in plant roots during watered conditions and not an enrichment of *Streptomyces* at under drought conditions. (**E**) Addition of an iron chelator to soil (ferrozine), results in elevated enrichment of *Streptomyces* in plant roots but not soil during drought (*n* = 8). (**F**) Increased enrichment is caused by a depletion of *Streptomyces* in plant roots in the presence of an iron chelator under watered conditions. Significance in BCDEF determined by Wald tests with FDR adjusted p values.

We next exploited the fact that iron becomes less bioavailable at alkaline soil pH to define the role of plant iron starvation responses in shaping *Streptomyces* enrichment in roots during drought. We experimentally raised soil pH from 5.81 to 7.39 and 7.99 with increasing lime amendment^56,65–67^ (Figure S3F). This resulted in a stepwise increase in the enrichment of *Streptomyces* in roots, but not soil, during drought (Figure 3C). Consistent with our model, we found that increased enrichment in roots resulted from decreasing *Streptomyces* abundance under watered conditions, rather than increasing abundance during drought at alkaline versus acidic pH (Figure 3D).

We then added ferrozine—a strong iron chelator—to the same soil to further show that iron limitation caused the effects observed at higher pH. Ferrozine reduces iron availability without causing soil disturbances associated with increased pH (Figure S3G and S3H). The addition of ferrozine increased the enrichment of *Streptomyces* exclusively in plant roots due to a reduction in their abundance under watered conditions (Figure 3E and 3F), as observed at elevated pH. This suggests that plant iron uptake activity — which increases at elevated pH and in the presence of an iron chelator but is suppressed during drought — inhibits *Streptomyces* accumulation.

Arabidopsis and other plants using the type I iron acquisition strategy produce phenolic compounds termed coumarins^52,68^, which help mobilize and can also reduce ferric iron from soil under iron limitation. Coumarins also have selective antimicrobial activity ^54–56^. We tested the inhibitory activity of fraxetin, one of the Arabidopsis coumarins produced under iron limitation at near neutral-alkaline conditions, against representative ASV1 strains spanning our collected soils and observed general inhibitory effects (Figure S3I), again consistent with our hypothesis. Collectively, these chemical manipulations suggest that rather than positively selecting *Streptomyces*, drought suppresses host plant immunity and iron uptake activity, which inhibit the proliferation of *Streptomyces* in plant roots (Figure 2G).

### Drought suppresses key components of the plant iron uptake apparatus

Antagonism between plant drought and immune responses is established^44–46,69^. We therefore investigated our observation that drought suppresses iron-responsive genes. Using a recently characterized “low-water” agar system^70^ (hereafter LWA, Table S5), in the absence of microbes, we found that at replete iron concentrations (+Fe) plants had reduced biomass relative to control conditions (MS) but no obvious signs of chlorosis (Figure 4A and 4B), a hallmark of iron starvation. However, at limiting iron concentration (-Fe), plants were severely chlorotic on LWA but not MS (Figure 4A and 4B). This LWA-induced chlorosis was not observed when we used other methods to impose osmotic stress on agar plates, including NaCl and polyethylene glycol (Figure S4A). The impact of LWA on chlorophyll production is dose dependent and reflects the combined effects of increased agar content and nutrient concentration (Figure S4e and S4F). LWA results in a root transcriptional response (performed independently^70^) that was largely shared among the shell and core DRGs from our wild soil dataset (Figure S4B). Thus, plant responses to LWA recapitulate the common DRGs observed in natural soil, including the finding that plants suppress iron-responsive genes resulting in pronounced chlorosis, making LWA a reasonable proxy for drought.

**Figure 4.**
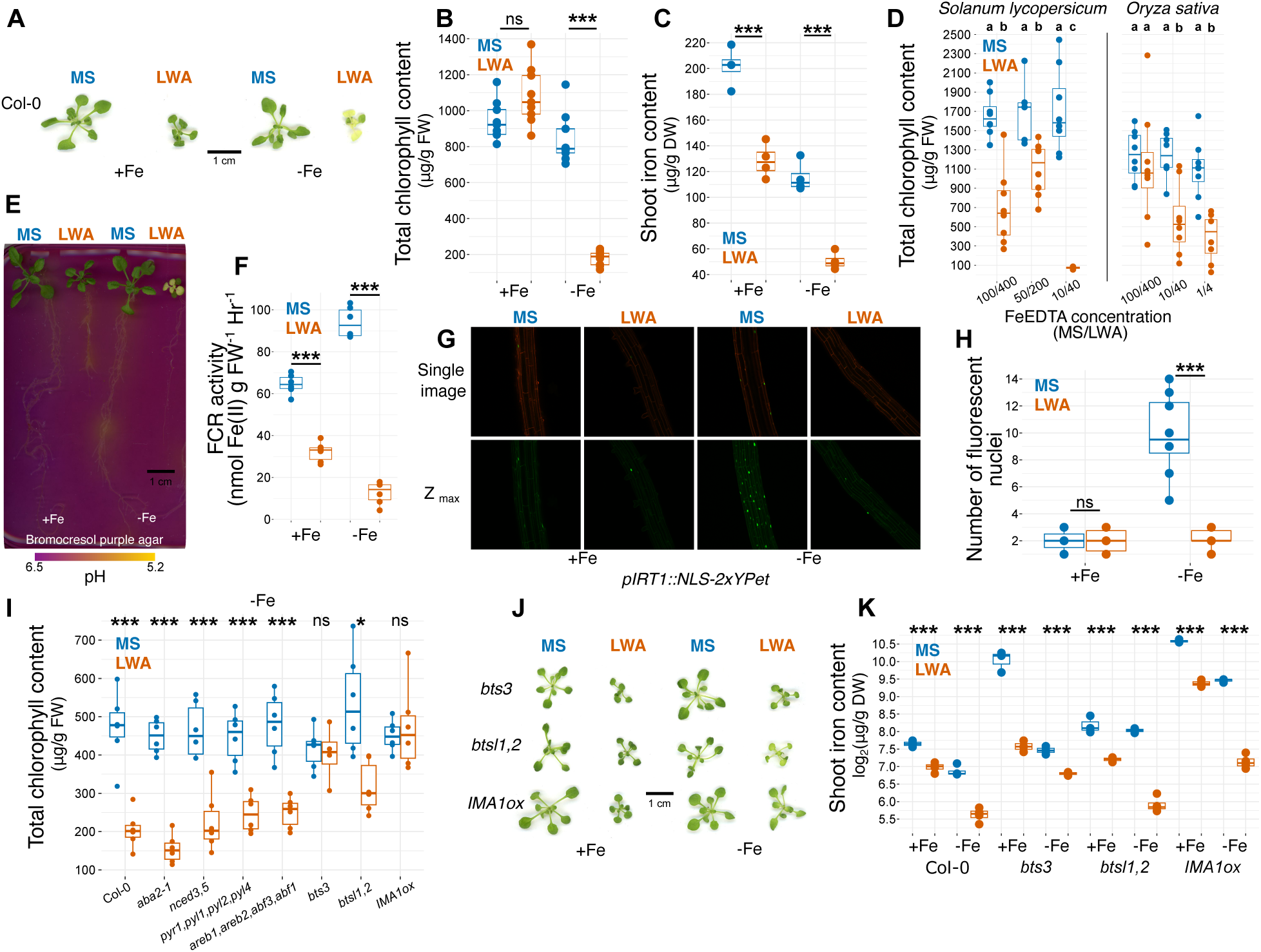
Drought induced suppression of plant iron uptake is independent of canonical water stress pathways. (**A**) Representative images of Col-0 plants grown on control (MS) or drought-mimicking media (LWA) at replete (+Fe) and limiting (-Fe) iron concentration. (**B**) Total chlorophyll (*n* = 9) and (**C**) iron content (*n* = 3) of Col-0 plants. (**D**) *Solanum lycopersicum* ‘micro-tom’ and *Oryza sativa* ssp. *japonica* ‘Nipponbare’ (*n* =6-8). Plants grown on LWA media exhibit suppression of (**E**) rhizosphere acidification (*n* = 6), (**F**) root ferric chelate reductase activity (*n* = 6), and (**G**) IRT1 promoter activity in roots. (**H**) Number of fluorescent nuclei of the pIRT1 reporter line grown (*n* = 6-8). (**I**) Total chlorophyll content of Col-0 and mutants perturbed in the ABA pathway or iron uptake (*n* = 6). (**J**) Representative images and (**K**) shoot iron content (*n* = 3) of mutants perturbed in iron uptake. For all experiments, plants were grown on 0.5X MS media for 7 days before transfer to experimental media and grown for 12 days (two-tailed t-test). Data in BCEGHJ were analyzed with two-tail t-tests.

Shoot iron concentration, a more direct readout of iron uptake, revealed a strong effect of LWA even under +Fe conditions (Figure 4C). Together with the chlorosis phenotype (Figure 4A), demonstrates that chlorophyll production is maintained when a threshold level of shoot iron concentration is met. We also measured shoot iron content of Col-0 growing in CHAP soil and found a large decrease in iron concentration during drought (Figure S4C). This confirmed the patterns of our transcriptional data and the observed effects of LWA on iron uptake. To test the generality of LWA-induced suppression of iron uptake, we grew *Solanum lycopersicum* (tomato) and *Oryza sativa* (rice), which span 160 Mya of plant diversity, on our LWA under varying levels of iron limitation. Like Arabidopsis, *S. lycopersicum* and *O. sativa* exhibited impaired iron uptake under -Fe conditions only on LWA (Figure 4D). We measured shoot iron content of *Triticum aestivium, Zea mays, Brassica napus,* and *Pisum sativum* grown in soil and observed a consistent reduction under drought (Figure S4D). These results reveal a conserved drought-induced suppression of iron uptake across plant species.

During iron limitation, iron acquisition strategy I plants such as Arabidopsis acidify their rhizosphere^71^ and upregulate key machinery of the reductive iron import apparatus, which includes the ferric reduction oxidase enzyme^71,72^, FRO2, and dedicated iron transporter^73^, IRT1. Plants on MS -Fe mounted an iron starvation response, as observed by rhizosphere acidification, increased ferric reductase activity and increased activity of the *IRT1* promoter (Figure 4E,4F,4G, and 4H). On LWA, all these inducible responses to -Fe were suppressed (Figure 4E,4F,4G, and 4H). These results demonstrate that LWA recapitulates the suppression of iron uptake observed during drought in wild soil and is generalizable across the monocot-eudicot divergence, encompassing species that engage in iron acquisition by either strategy I or II.

To understand how the suppression of iron uptake is integrated into existing drought response pathways, we grew ABA mutants impaired in the biosynthesis^74^ (*nced3 nced5* and *aba2*) perception^75^ (*pyr1 pyl1 pyl2 pyl4*), and transcriptional responses^76^ (*areb1 areb2 abf3 abf1*) on LWA-Fe. LWA-induced suppression of iron uptake was entirely ABA independent (Figure 4I), in line with our microbiome community analyses demonstrating that *Streptomyces* enrichment during drought is also ABA independent (Figure S2C). We also found no role for the dehydration responsive element^77^ (DRE: *drip1 drip2*) pathway in the modulation of iron uptake ability (Figure S4G). These results indicate that the suppression of iron uptake on LWA occurs independently of the canonical drought response pathways (which were also defined using simplified growth systems).

We next asked whether known regulatory elements in the iron starvation response might be required for our observed chlorosis on LWA-Fe. Several RING E3 ligases including BRUTUS (BTS) and BRUTUS LIKE1 (BTSL1) and BRUTUS LIKE2 (BTSL2) negatively regulate iron starvation responses by targeting key transcription factors for proteasomal degradation^78,79^. We assessed whether known regulatory elements in the iron starvation response are required for our observed chlorosis on LWA-Fe. *bts-3* exhibited a complete restoration of chlorophyll production on LWA-Fe, while the double mutant *btsl1 btsl2* had a partial restoration (Figure 4I and 4J). We also tested transgenic plants overexpressing IMA1, a small peptide and positive regulator of iron starvation responses^80,81^. We observed restoration of chlorophyll production (Figure 4I and 4J). IMA1 sequesters BTS and thus reduces the ubiquitination of its primary targets^78^, basic helix–loop–helix subgroup IVc transcription factors 105 and 115^79^, which positively regulate the expression of iron responsive genes. However, like Col-0, when we measured shoot iron concentration, *bts-3*, *btsl1 btsl2,* and *IMA1ox*, all exhibited a reduction in iron concentration on LWA, regardless of iron status (Figure 4K). Given our previous finding that chlorophyll production is maintained when a threshold of shoot iron concentration is met, it is likely that LWA-induced suppression of iron uptake is masked in these mutants even under -Fe because of their ability to hyperaccumulate iron. Collectively, these results suggest that LWA-induced suppression of iron uptake is orchestrated by negative regulation of the iron starvation response, which occurs upstream of IMA1 and independently of the canonical water stress response governed by the ABA and DRE pathways. The suppression of the iron starvation response during drought has a clear impact on the plant and, as our findings demonstrate, also influences the root enrichment of *Streptomyces*.

### Plant benefit provisioned by *Streptomyces* isolates requires a functioning reductive import apparatus and suppression of plant immunity

The enrichment of particular plant-associated microbes under stress is often interpreted as a beneficial response to a plant ‘cry for help’ response^23,24^. We demonstrated that suppression of SA immunity and iron uptake contribute to the drought enrichment of *Streptomyces* in roots, but the functional consequences of *Streptomyces* enrichment remain unclear. We tested whether microbes had the capacity to rescue either the biomass cost of LWA in iron replete conditions (MS+Fe vs. LWA+Fe) or the chlorophyll cost of LWA in the presence in iron limiting conditions (MS-Fe vs. LWA-Fe) using the axenic system (Figure 5A).

**Figure 5.**
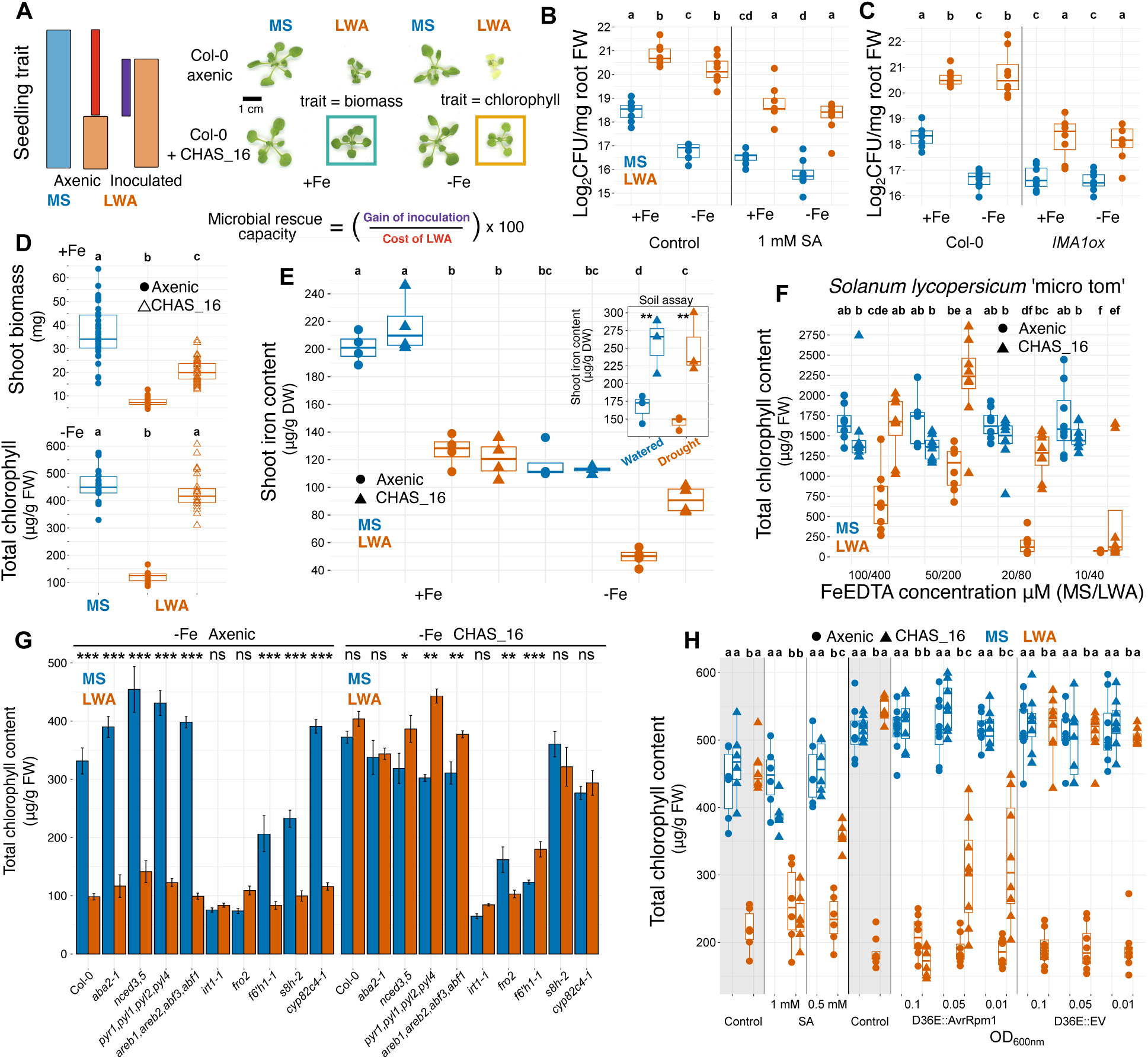
Benefit provisioned to plants by *Streptomyces* depends on the iron reductive import system in plants and the suppression of plant immunity. (**A**) Rescue capacity is the proportion of the cost of LWA under axenic conditions, measured as either biomass or chlorophyll reduction, recovered by microbial inoculation. (**B**) CFU counts of CHAS_16 from Col-0 roots (*n* = 8). (**C**) CFU counts of CHAS_16 from *IMA1ox* roots (*n* = 8). (**D**) Quantification of the CHAS_16 biomass and chlorophyll rescue capacity (*n* = 18). (**E**) Shoot iron content of Col-0 grown axenically or with CHAS_16 in agar or soil (inset, *n* = 3-4). (**F**) CHAS_16 rescues chlorophyll production in the ‘micro-tom’ *Solanum lycopersicum* genotype (*n* = 8). (**G**) The Col-0 genetic requirements of chlorophyll rescue by CHAS_16 (*n* = 6). The left panel shows the effects of LWA-Fe under axenic conditions and the right, the same effects in the presence of CHAS_16. (**H**) The chlorophyll rescue provisioned by CHAS_16 on LWA-Fe is abolished when plant immunity is activated with exogenous SA application or inoculation with *Pseudomonas syringae* pv. tomato DC3000 D36E::AvrRpm1 (*n* = 5-6). Data in BCDEFH were analyzed using ANOVA followed by Tukey’s test where letters indicate statistically significant differences between different conditions. Data in G were analyzed with two-tail t-tests.

Using CHAS_16, an exemplary *Streptomyces* ASV1 isolate (Figure 5A and S5A), we confirmed in the agar-based system that root enrichment occurs on LWA versus MS. The magnitude of the enrichment of CHAS_16 increases when we impose iron limitation, due to a decrease in abundance on MS-Fe (Figure 5B and 5C). When inoculated on the iron hyperaccumulator *IMA1ox*^80,82^, CHAS_16 reached lower abundance on both MS and LWA, as anticipated (Figure 5C). As observed in CHAP soil, CHAS_16 abundance decreased when plants were treated with an immune activator, SA (Figure 5B). To test whether immune activation under more natural conditions recapitulates the effects of exogenous SA, we inoculated plants with the well-studied foliar pathogen *Pseudomonas syringae* pv. *tomato* DC3000 (PtoDC3000)^83^. PtoDC3000 suppresses plant immunity via type III secretion system (T3SS)–delivered effector proteins. We compared a non-virulent mutant lacking all 36 effectors (D36E) with D36E::AvrRpm1, in which the ETI-eliciting effector AvrRpm1 is restored^84,85^. Consistent with SA treatment, D36E:AvrRpm1 reduced CHAS_16 abundance in roots, while the non ETI-eliciting strain D36E::EV did not (Figure S5B).

On LWA, CHAS_16 increased plant biomass under +Fe conditions and maintained chlorophyll production under -Fe conditions (Figure 5D). Chlorophyll rescue required a live bacterium and was stable across two orders of magnitude of bacterial density (Figure S5C and S5D). CHAS_16 maintained rhizosphere acidification and increased IRT1 promoter activity on LWA–Fe, indicating that chlorophyll rescue is accompanied by enhanced plant iron starvation responses (Figure S5E and S5F). CHAS_16 also maintained shoot iron content of plants grown on LWA–Fe (Figure 5E) and increased iron content of *Arabidopsis* grown in an inert soil substrate under both watered and drought conditions (Figure 5E inset).

To test whether this rescue extends beyond *Arabidopsis*, we inoculated *S. lycopersicum* grown on LWA across a range of iron concentrations. CHAS_16 restored chlorophyll production at all but the most limiting iron levels (Figure 5F). A similar limitation was observed in *Arabidopsis* (Figure S5G), where CHAS_16 maintained chlorophyll production relative to axenic plants grown on MS but not on LWA under extremely low Fe. Iron limitation can also occur at high pH^52^, especially when iron is supplied as less bioavailable FeCl_3_^54,86^, we tested these conditions and found that CHAS_16 restored chlorophyll production across iron concentrations and forms at alkaline pH (Figure S5H).

We found that CHAS_16 chlorophyll rescue capacity was entirely ABA independent (Figure 5G) but required an intact iron reductive import system (IRT1 and FRO2), like the endogenous suppression of iron uptake during drought. We tested four additional *Streptomyces* isolates with high chlorophyll rescue capacity and found identical plant genetic requirements, suggesting that among beneficial *Streptomyces* isolates, a common mechanism exists to maintain plant iron uptake during drought (Figure S5I). In contrast, none of the tested candidate plant genes were required for biomass rescue capacity on LWA+Fe (Figure S5J), indicating that biomass and chlorophyll rescue occur via independent plant-encoded pathways.

Through experiments where plants were treated with SA or PtoDC3000 D36E::AvrRpm1, we showed that the chlorophyll rescue capacity of CHAS_16 was attenuated by dose-dependent immune activation (Figure 5H). Treating plants with 1mM SA or 0.1 OD_600_ PtoDC3000 D36E::AvrRpm1led to the complete loss of the CHAS_16 chlorophyll rescue capacity. This indicates that the maintenance of plant immune activation on LWA, either via SA application or PtoDC3000 D36E::AvrRpm1 infection, desensitizes plants to the benefit provisioned by CHAS_16. We repeated the experiment using three additional *Streptomyces* isolates with chlorophyll rescue capacity (Figure S5K) and found that treatment with 1 mM SA consistently eliminated this benefit.

Given the reported widespread ability of diverse bacteria to rescue chlorophyll production during pH mediated iron starvation^54^, we investigated how general this chlorophyll-based rescue capacity is on LWA-Fe. We tested 30 non-*Streptomyces* bacteria that represent core taxa within the Arabidopsis root microbiome and found only one strain that could rescue chlorophyll production on LWA-Fe (Figure S5L). Thus, the benefit provisioned to plants by CHAS_16, which includes the maintenance of iron uptake, occurs across diverse conditions and represents a trait enriched among *Streptomyces*.

### Plant benefit provisioned by *Streptomyces* is not related to enrichment and is modulated by intra-genus antagonism

Our data shows that in both watered and drought conditions roots are colonized by consortia of *Streptomyces.* To mimic these consortia, we assembled 10 synthetic communities (SynCom) comprising isolates that matched the sequence of enriched (E) and non-enriched (NE) *Streptomyces* ASVs. These SynCom capture much of the *Streptomyces* community in plant roots from 10 soils spanning the observed variation in enrichment (Figure 6A, S6A, and S6B). Surprisingly, SynCom varied widely in their capacity to rescue biomass or chlorophyll on LWA, with no correlation to the observed *Streptomyces* enrichment in corresponding soils (Figure 6A). We decomposed SynCom into individual E and NE isolates and measured their rescue capacity in mono-association. Again, we observed wide variation in biomass and chlorophyll rescue capacity, independent of enrichment status and soil location (Figure 6A). This demonstrates that enrichment status, sequence identity or soil origin do not predict a *Streptomyces* isolate’s rescue ability.

**Figure 6.**
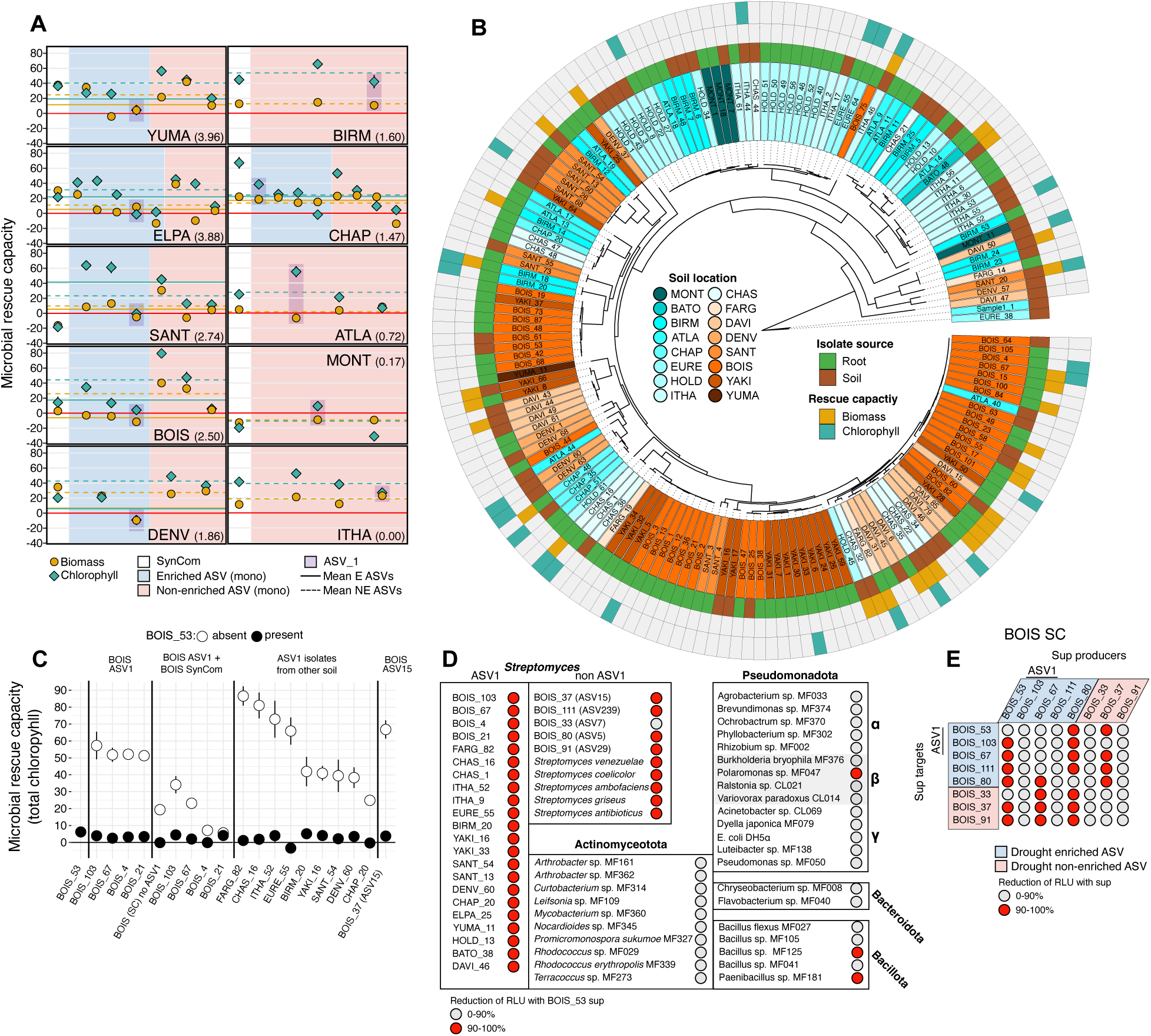
Benefit provisioned to plants by *Streptomyces* is decoupled from enrichment status and is strongly modulated by intra-genus antagonism. (**A**) Rescue capacity of small, soil-specific synthetic communities (SynComs; SC; white backgrounds) and individual SC members corresponding to isolates matching enriched (E; blue background) or non-enriched (NE; red background) ASV sequences. Both biomass (LWA+Fe, yellow symbols/lines, *n* = 6-36) and chlorophyll (LWA-Fe, green symbols/lines, *n* = 6-12) rescue are shown. Mean values represented with lines (E strains: solid line; NE strains: dashed line). Panels are ordered by the magnitude of *Streptomyces* enrichment, stated next to the soil location. The rescue capacity of ASV1 is highlighted with a purple box. (**B**) A whole genome phylogeny of ASV1 isolates coming from 16 soils. The innermost ring depicts soil location, followed by isolate source (cultured from roots or soil), biomass rescue capacity, and chlorophyll rescue capacity. (**C**) BOIS_53 eliminates the chlorophyll rescue capacity of diverse single strains and consortia of *Streptomyces* (*n* = 6). (**D**) Liquid culture supernatant derived from BOIS_53 exhibits strong inhibitory activity confined to targets within *Streptomyces* (*n* = 3-6). (**E**) Reciprocal supernatant inhibition assays among BOIS SC members reveals stronger inhibitory activity between E producers and NE targets versus NE producers and E targets (*n* = 3-6).

Isolates matching the widely enriched ASV1 exemplified this pattern (Figure 6A). Using a large ASV1 isolate collection derived from roots of plants grown in our diverse soils and matching unplanted soil under drought conditions, we quantified rescue capacity and performed whole genome sequencing of 164 ASV1 isolates from 16 soils. ASV1 isolates exhibited a high degree of diversity at the genomic level and in their rescue capacity, the extent of which was largely recapitulated at each soil location (Figure 6B). Given that ASV1 isolates were often cultured from the same root or soil sample, and thus co-occur, we investigated the rescue outcome when isolates with divergent phenotypic effects were co-inoculated with plants under LWA-Fe. We focused on BOIS isolates because of the number of ASV1 isolates from roots grown in this soil, the strong enrichment of BOIS ASV1 during drought, and their span of chlorophyll rescue capacity (Figure 6B). First, we inoculated plants with BOIS ASV1 isolates with high chlorophyll rescue capacity in the presence or absence of BOIS_53, an ASV1 isolate initially used for the SC experiments that demonstrated poor rescue capacity. The rescue capacity of all BOIS ASV1 strains was eliminated in the presence of BOIS_53 (Figure 6C). Next, we observed that replacing BOIS_53 with other BOIS ASV1 isolates significantly increased the chlorophyll rescue capacity of the BOIS SC as a whole and that these gains were eliminated when co-inoculated with BOIS_53 (Figure 6C). Co-inoculation with BOIS_53 eliminated the chlorophyll rescue capacity of ASV1 strains isolated from nine additional soil locations (Figure 6C). BOIS_53 also eliminated the benefit of BOIS_37, a non-ASV1 strain with high chlorophyll rescue capacity, even at a 10-fold lower abundance (Figure S6D). Collectively, these results highlight the sub-ASV diversity in the potential to provision a benefit to the plant among ASV1 strains isolated from the same source material. These results also caution against any general functional attribution, beneficial or otherwise, to at least this clade based on its prevalent enrichment in plant roots during drought.

We hypothesized that the dominant negative effect of BOIS_53 on the chlorophyll rescue across diverse *Streptomyces* was due to growth inhibition via secreted toxins. *Streptomyces* produce highly specific polymorphic toxins termed umbrella particles^87^, which inhibit growth and mediate competition among close relatives. We prepared BOIS_53 supernatant, which excludes small molecules (>100 kDa; sup*BOIS_53*), and tested its activity against ASV1 isolates whose rescue capacity was masked by BOIS_53, as well as additional ASV1 isolates from each soil. In contrast to the specific inhibitory effects of umbrella particles, growth of every tested isolate was severely inhibited when supplemented with sup*BOIS_53* (Figure 6D) as was the chlorophyll rescue capacity of BOIS_37 *in planta* when supplemented with sup*BOIS_53* (Figure S6D). We expanded the target strains to include all members of the BOIS SynCom, commonly used model *Streptomyces* strains, and diverse bacterial isolates representing core members of the root microbiome in Arabidopsis. We observed a remarkable breadth of sup*BOIS_53* inhibitory activity within *Streptomyces*, but high specificity to strains from this genus since inhibition in non-*Streptomyces* was rare (Figure 6D). These results demonstrate that BOIS_53, a representative isolate of the highly REC enriched ASV1, eliminates the plant benefits provisioned by other *Streptomyces*, including those enriched in the REC during drought, via secreted proteins that inhibit the growth of congeneric competitors.

We performed a fully reciprocal supernatant inhibition assay among representative isolates matching either E or NE ASVs from BOIS to understand how intra-*Streptomyces* antagonism might contribute to the observed relative drought enrichment among *Streptomyces* ASVs (Figure 6E). Like BOIS_53, supernatant from BOIS_80, which matched the sequence of an E ASV, exhibited broad inhibitory activity against every BOIS SynCom target, regardless of enrichment status (Figure 6E). The supernatant derived from NE strains had little inhibitory activity, except BOIS_37 (Figure 6E). We generated supernatant from BOIS_103 and BOIS_67, both of which have chlorophyll rescue capacity and are inhibited by sup*BOIS_53*, to test whether the broad inhibitory activity observed with sup*BOIS_53* was conserved among BOIS ASV1 isolates. sup*BOIS_103* showed no activity against any BOIS SynCom target, while sup*BOIS_67* had activity primarily against NE strains and thus dissimilar from the broad activity of sup*BOIS_53* (Figure 6E). These results suggest that intra-genus competition could drive relative enrichment among co-occurring *Streptomyces* in roots during drought. Combined with our finding that enrichment status is unlinked to benefit provisioning, this suggests that the *Streptomyces* traits responsible for plant benefit evolved independently from those contributing to relative enrichment in plant roots, the latter being dictated by interactions with congeneric bacteria. Such interactions affect microbiome community assembly but can also be consequential for plant host health, as shown by the dramatic elimination of rescue capacity by BOIS_53 across a range of community contexts. Thus, under drought, *Streptomyces* exploit plant roots as a niche, where competitive success does not guarantee benefits to the plant.

## Discussion

When plants migrated to the terrestrial environment 450 MY ago, they encountered novel stressors including periods of water scarcity^88,89^. In response, plants evolved morphological and anatomical traits that reduced water loss as well as physiological responses that limited the negative effects of drought^90–95^. Throughout, plants were in close association with microbes, thus, it is plausible that plant responses to environmental stress are integrated with selection for some or all of the resident microbiome^96,97^. The enrichment of *Streptomyces* in plant roots during drought is a widely distributed plant microbiome phenotype occurring across diverse plant taxa^29–32^. We add to this overall generality by demonstrating that enrichment on Arabidopsis roots is remarkably prevalent across diverse natural soils with high temporal consistency (Figure 1), suggesting that the enrichment capacity is geographically widespread among *Streptomyces*.

The widespread enrichment of *Streptomyces* in roots during drought led to the idea that this phenotype reflects a plant ‘cry for help’ to alleviate drought stress. Our findings refine this view, suggesting instead that *Streptomyces* exploit the plant root as a conditional niche during drought because host suppression of plant immunity and iron homeostasis removes otherwise inhibitory activities (Figures 2–4). Among *Streptomyces* strains, enrichment during drought likely reflects competitive interactions governed by polymorphic secreted protein toxins. Competitive success and the ability to provide plant benefits under drought, including biomass production and iron uptake, are unlinked (Figure 6); thus enrichment does not guarantee plant benefit. Similar decoupling of stress-induced microbial enrichment and plant benefit occurs across plant lineages under diverse stresses^54,98,99^.

Plant root and shoot transcriptional profiling revealed that despite a core drought response, differential gene expression varies widely across soils and correlates with soil and environmental attributes (Figures 2 and S2). The functional importance of these associations remains unclear but will likely be important for managing plant growth during drought across diverse soils. We discovered that expression of SA- and iron-responsive genes covaries in roots across natural soils (Figure 2). Genetic or physiological manipulation of these processes directly modulates *Streptomyces* abundance under watered conditions and consequently alters the magnitude of enrichment during drought through suppression of plant immunity and iron homeostasis (Figures 3 and 4). Plant immune and physiological activity, particularly nutrient homeostasis, are established drivers of plant microbiome composition and diversity^12–14,50,54^. Thus, soil locality shapes the plant microbiome not only by providing the source of microbial colonists but also by influencing plant physiological and immune processes.

Suppression of immunity during drought occurs across plant species and is thought to arise from antagonism between ABA and SA signalling and prioritization of abiotic stress responses over immune activation^46^. In contrast, suppression of iron uptake during drought remains less explored^100–102^. Excess iron can promote the formation of reactive oxygen species^53^, suggesting plants might restrict iron uptake during periods of reduced growth to avoid toxicity. However, under osmotic stress, iron hyper-accumulating mutants produced more biomass and maintained chlorophyll production relative to Col-0 (Figure 4), suggesting this interpretation should be reconsidered and highlighting iron homeostasis as a potential target for improving drought resilience. This is particularly important given the widespread prevalence of iron-deficient soils, the reliance on plants as major dietary iron sources, and increasing aridity across agricultural regions. Drought-induced suppression of plant iron uptake, together with reduced baseline immune activity, likely creates conditions that enable *Streptomyces* enrichment across diverse land plants. Why this suppression benefits *Streptomyces* remains unclear, although the coincidence of plant terrestrialization and *Streptomyces* diversification suggests that drought-associated colonization of plant roots may have influenced the clade’s evolution.

Our functional analyses did not support the hypothesis that enrichment of *Streptomyces* during drought is necessarily linked to plant benefit at the level of consortia or individual strains (Figure 6). The ability to enhance plant growth or maintain chlorophyll production under drought-like conditions was rare among isolates of the cosmopolitan and widely enriched ASV1 and also occurred among strains isolated from soils showing minimal enrichment (Figure 6). *Streptomyces* possess traits that could benefit plants under drought and iron limitation^103,104^, including exploratory growth^105,106^, and production of pteridic acid^107^. For CHAS_16, an exemplary ASV1 isolate with plant rescue ability, chlorophyll rescue in *Arabidopsis* required an intact host iron reductive-import system, suppression of SA-triggered immunity, and enhancement of the endogenous iron starvation response (Figures 5 and S5). Identifying the genetic mechanisms that enable plants to receive, and bacteria to deliver, enhanced biomass and chlorophyll production during drought will be an important direction for future work.

Co-inoculation experiments revealed that competitive interference mediated by secreted high-molecular-weight proteins can eliminate benefits provided by individual strains in pairwise and community contexts (Figure 6C). One isolate, BOIS_53, showed broadly inhibitory activity restricted to *Streptomyces* (Figure 6D). This pattern differs from recently described umbrella toxins^87^ and warrants further investigation. Differential inhibition among enriched and non-enriched isolates from BOIS may help explain enrichment patterns (Figure 6E). Similar competition among strains differing in plant benefit occurs in established symbioses such as legumes–rhizobia^108,109^ and plants–mycorrhizae^110,111^. Thus, the presence of enriched strains that fail to confer benefit does not exclude potential advantages from broader *Streptomyces* enrichment within the REC. However, our finding that enrichment status is uncoupled from plant benefit (Figure 6A) suggests that traits underlying plant benefit evolved independently from those promoting proliferation in plant roots during drought. This decoupling challenges a simple ‘cry for help’ model. Instead, drought-induced suppression of host iron uptake and immunity promotes *Streptomyces* enrichment, while intra-genus interactions refine strain-level composition and functional outcomes.

### Limitations of the study

In this study, we demonstrated widespread *Streptomyces* enrichment in roots of a single plant genotype during drought in natural soils under controlled growth conditions. This design enabled comparisons across soils while maintaining natural soil contexts. Including additional plant species will help determine the generality of these findings. We show that drought suppresses plant iron uptake, identify *Streptomyces* capable of maintaining iron uptake under drought-like conditions, and uncover widespread inhibitory interactions among root-associated *Streptomyces*. However, the genetic mechanisms underlying these processes remain unresolved and represent important targets for future work.

## Resource availability Lead contact

Requests for further information and resources should be directed to and will be fulfilled by the lead contact, Jeff Dangl.

## Materials availability

Bacterial isolate strains and plant materials in this study will be made available on request to the lead contact, Jeff Dangl.

## Data and code availability

The bacterial 16S rRNA, RNA-seq, and whole genome sequence files have been deposited in the National Center for Biotechnology Information (NCBI) SRA database (16S rRNA and WGS: PRJNA1285979, RNA-seq: PRJNA1287558). Intermediate data and analysis scripts employed in the analyses are available on Mendeley data: https://data.mendeley.com/preview/mz2kb4ych5?a=99697200-5370-496f-a412-c9775a245a38.

## Acknowledgments

We thank all the personnel at the local organizations who collected and shipped soil. We thank members of the UNC high-throughput sequencing facility for their service and support throughput the project. We thank T. Interrante, J. Garzoni and J. Winshell for technical assistance and members of the Kieber lab (University of North Carolina at Chapel Hill, USA) and He lab (Duke University, USA) for helpful discussion. This work was supported by NSF grant IOS-1917270 to J.L.D and C.D.J. J.L.D. is an HHMI Investigator and John N. Couch endowed Professor. J.D.M. is an HHMI Investigator and the Lynn M. and Michael D. Garvey Endowed Chair in Gastroenterology. C.R.F gratefully acknowledges funding by a Natural Sciences and Engineering Research Council of Canada postdoctoral fellowship (532852-2019). D.R gratefully acknowledges funding by EMBO Long Term Fellowship (ALTF 743–2019). We thank Drs. O. Finkel, K. Paul, and P.J.P.L Teixeira for critical comments and the Dangl lab microbiome team for comments throughout the course of this project. This article is subject to HHMI’s Open Access to Publications policy. HHMI lab heads have previously granted a nonexclusive CC BY 4.0 license to the public and a sublicensable license to HHMI in their research articles. Pursuant to those licenses, the author accepted manuscript of this article can be made freely available under a CC BY4.0 license immediately upon publication.

## Author contributions

C.R.F. and J.L.D. conceived the study. J.L.D, S.R.G., and C.D.J. supervised the project. C.R.F., R.A.S, J.H., T.F.L., D.R., A.A.E., P.J., N.K., C.S., M.A.M., O.E.A. performed the experiments. R.A.S. and C.D.J. analyzed the RNAseq data and W.Z. and C.D.J. analyzed the genomic data. M.J. and C.T.U.L. assisted with experiments. T.S. assisted with genomic library prep and sequencing. Q.Z. generated the *Streptomyces* supernatant and J.D.M., Q.Z., and S.B.P. guided the inhibition experiments. C.R.F., C.D.J., and J.L.D. wrote the manuscript with input from all authors.

## Declaration of interests

The authors declare no competing interests.

## Supplemental information

**Figure S1.**
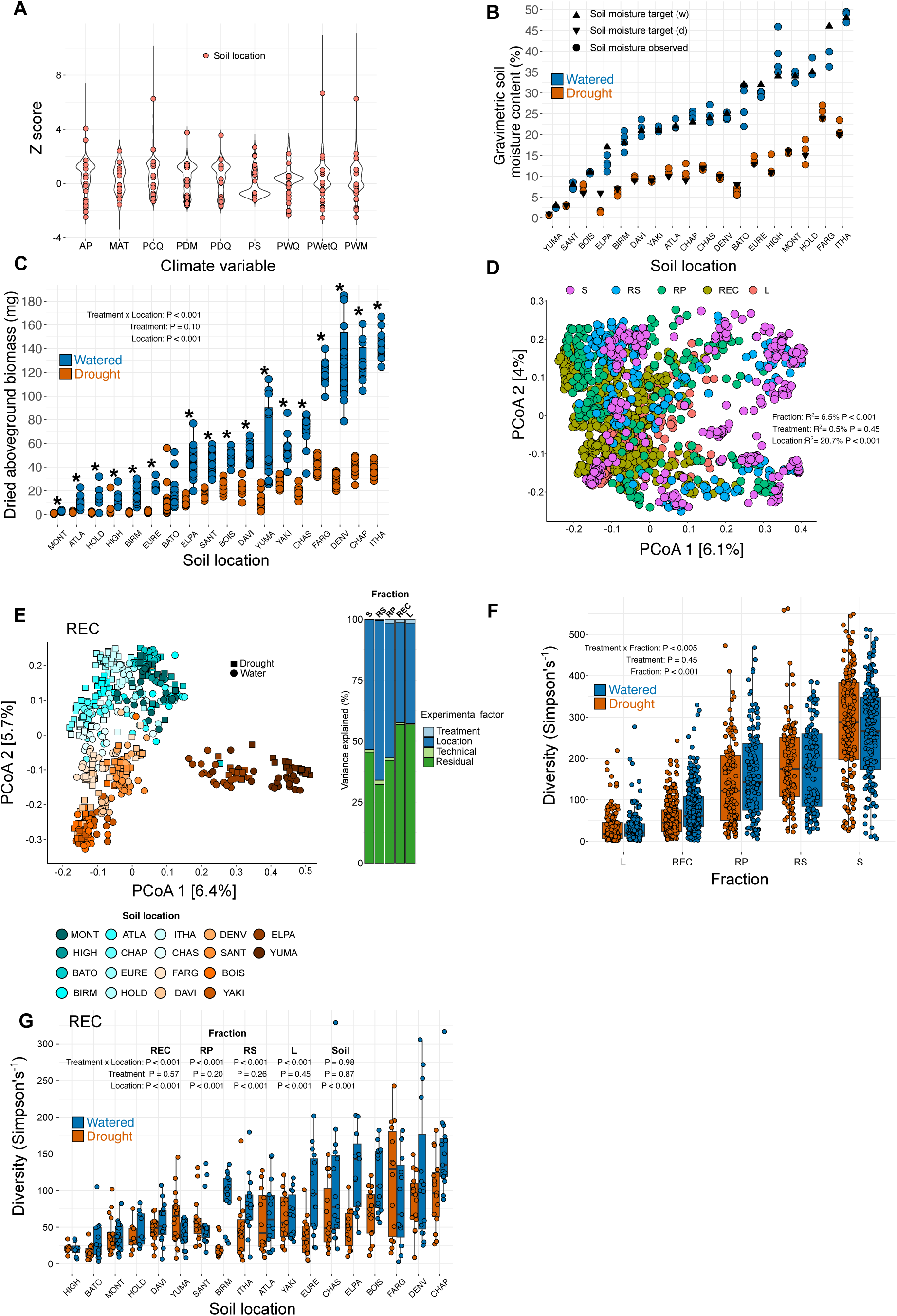
The effects of drought on the diversity and composition of plant microbiome habitats, Related to Figure 1. (**A**) Soil locations (points) were selected to span climatic variation (violin plot) in the contiguous USA. Each violin plot shows the kernel probability density of the values of 28,838 data points across the contiguous USA. Climate variables: AP annual precipitation, MAT mean annual temperature, PCQ precipitation in coldest quarter, PDM precipitation in dryest month, PDQ precipitation in dryest quarter, PS precipitation seasonality, PWQ precipitation in warmest quarter, PWetQ precipitation in wettes quarter, PWM precipitation in wettest month. (**B**) Target and observed gravimetric soil moisture content to achieve our desired soil moisture states of field capacity and permanent wilting point (*n* = 3-4). (**C**) Dried aboveground biomass of Col-0 plants grown under watered and drought conditions in each soil (*n* = 6-15). (**D**) Principal coordinate analysis of Bray-Curtis dissimilarity among bacterial community samples (S: *n* = 317, RS: *n* = 228, RP: *n* = 240, REC: *n* = 518, L: *n* = 296). PERMANOVA results shown on figure. (**E**) Principal coordinate analysis of Bray-Curtis dissimilarity among REC bacterial community samples (*n* = 7-20). PERMANOVA results for the tested experimental factors shown on right side for each plant microbiome habitat. (**F**) Bacterial community diversity across habitats and watering treatment (S: soil, *n* = 317, RS: rhizosphere, *n* = 228, RP: rhizoplane, *n* = 240, REC: root endophytic compartment, *n* = 518, L: leaf, *n* = 296). (**G**) Bacterial diversity in the REC shown across locations and watering treatment (*n* = 7-20) with ANOVA results displayed for each habitat. Significance in C,F,G determined by ANOVA. Statistical significances indicated by ∗∗∗ p < 0.001, ∗∗ p < 0.01, ∗ p < 0.05. For C,F,G (and all subsequent boxplots presented) the horizontal bars within the boxes represent the medians. The tops and bottoms of the boxes represent the 75th and 25th percentiles, respectively. The upper and lower whiskers represent 1.5× the interquartile range from the upper edge and lower edge of the box.

**Figure S2.**
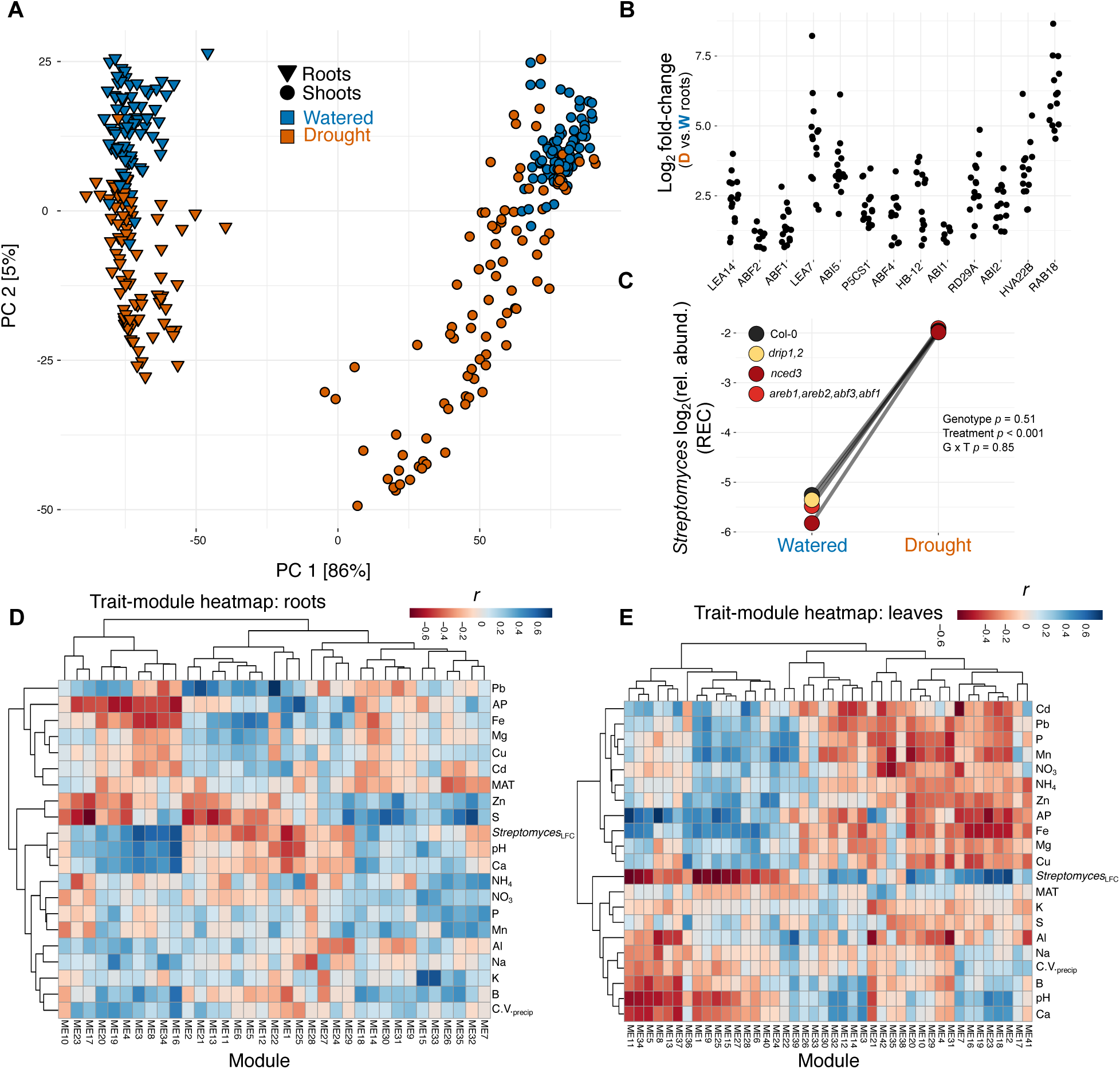
Core drought responses do not contribute to *Streptomyces* enrichment in roots during drought, Related to Figure 2. (**A**) Global Principal Component Analysis of the transcriptomes of leaves and roots among all 15 soils under watered and drought conditions. (**B**) ABA-responsive genes are among the core upregulated DRGs in roots. Each point represents the log_2_ fold-change (all *p_adj_* < 0.05) of a given gene at a one of the 15 soil locations. (**C**) Arabidopsis mutants perturbed in the canonical plant drought response pathways, ABA (*nced3, areb1 areb2 abf3 abf1*) and DRE (*drip1 drip2*), enrich *Streptomyces* in roots during drought (*n* = 8-10). (**D**) Full trait-module heatmap showing the correlations between the differential expression of gene modules (drought versus watered condition) defined in roots and soil attributes (*n* = 15). (**E**) Full trait-module heatmap showing the correlations between the differential expression of gene modules (drought versus watered condition) defined in leaves and soil attributes (*n* = 15). Significance in B,C determined by Wald tests with FDR adjusted p values.

**Figure S3.**
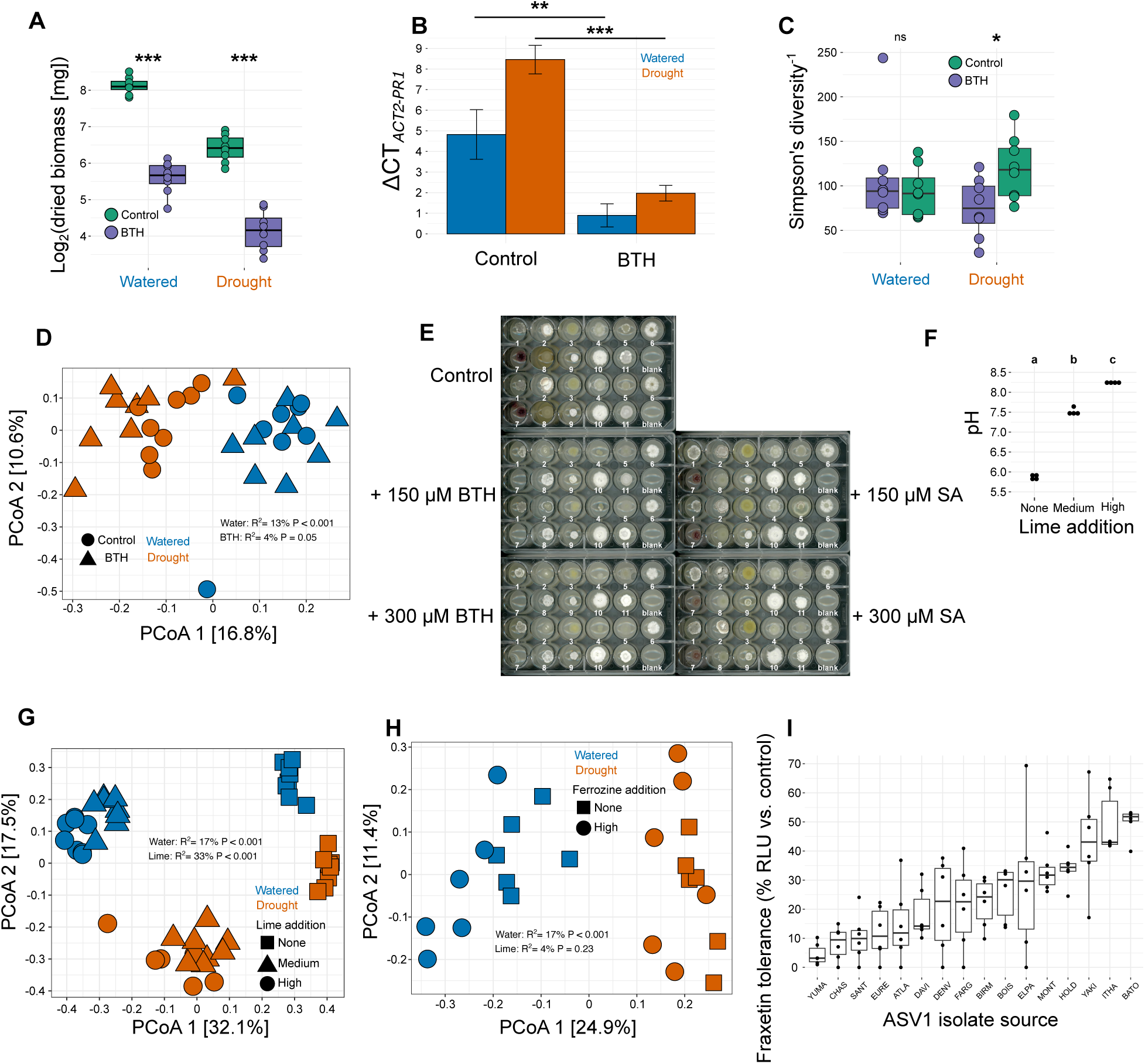
Manipulation of plant immunity and soil iron availability modulates *Streptomyces* abundance in roots, Related to Figure 3. (**A**) Dried aboveground biomass of plants across chemical and watering treatments (*n* = 8). (**B**) ΔCT values (*ACT2* – *PR1*) of the SA immune marker gene *PR1* in leaves across chemical and watering treatments (*n* = 8, error bars represent the standard error). (**C**) Bacterial diversity in the REC across chemical and watering treatments (*n* = 7-8). (**D**) Bacterial community composition in the REC across chemical and watering treatments (*n* = 7-8). (**E**) *Streptomyces* growth is largely unaffected by supplementation with BTH or SA. Each plate contains two replicates of 11 *Streptomyces* strains isolated from CHAP soil and includes representatives of all members of the CHAP SynCom. (**F**) pH of CHAP soil after lime amendment (*n* = 4). (**G**) Bacterial community composition in the REC across pH and watering treatments (*n* = 8). (**H**) Bacterial community composition in the REC with the addition of ferrozine during watered and drought conditions (*n* = 6). (**I**) Inhibitory effects of the plant coumarin fraxetin on *Streptomyces* ASV1 isolates from across soil localities (*n* = 6). Statistical significances indicated by ∗∗∗ p < 0.001, ∗∗ p < 0.01, ∗ p < 0.05.

**Figure S4.**
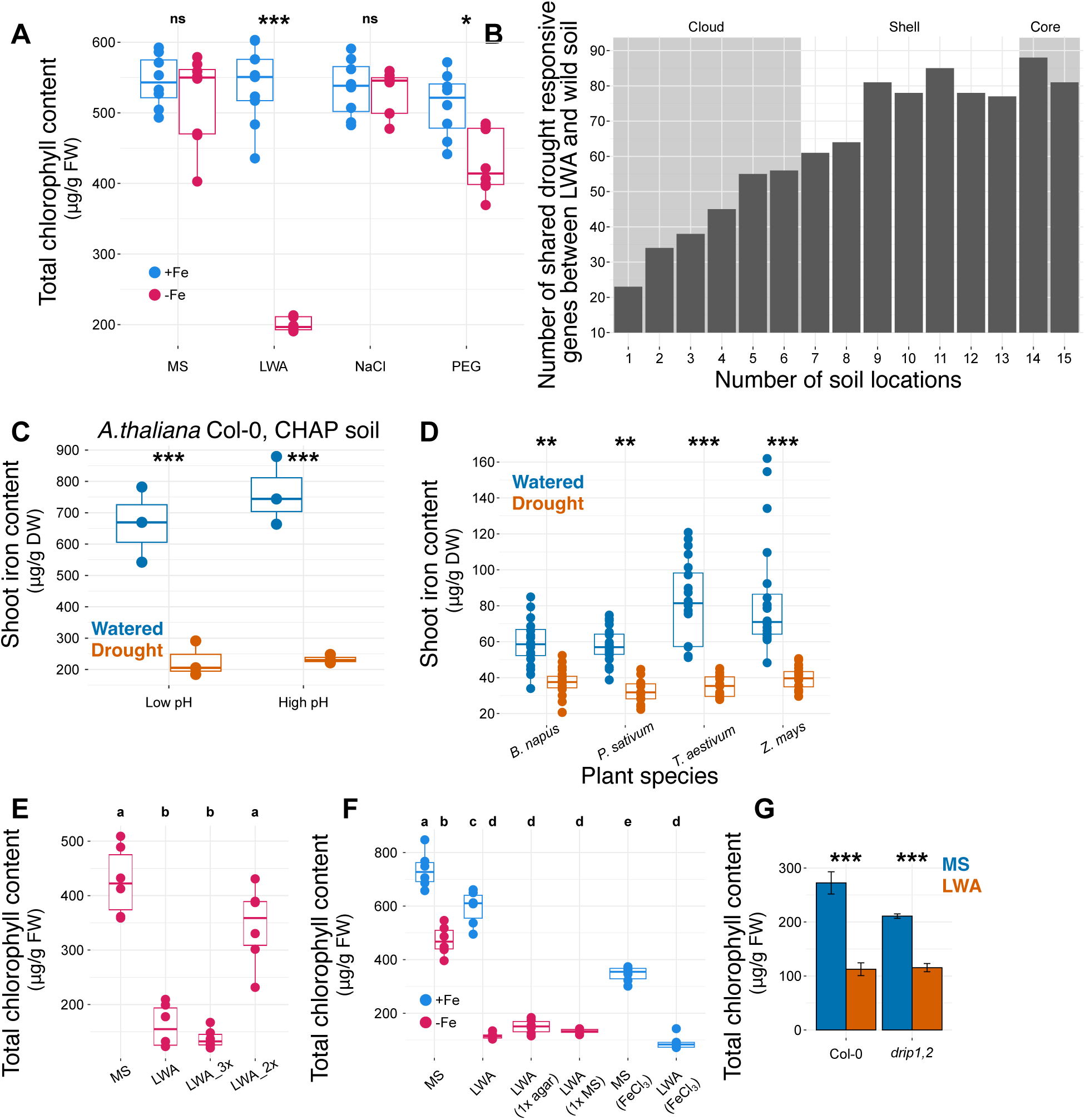
LWA recapitulates core drought-responsive genes across natural soils and results in suppression of iron uptake across plant species, Related to Figure 4. (**A**) Total chlorophyll content of Col-0 seedlings grown under replete (+Fe) and limiting (-Fe) iron conditions on different media types (*n* = 8, NaCl concentration 100 mM, PEG water potential -1.2 MPa). (**B**) The majority of differentially expressed genes in Col-0 roots in response to LWA were found in the core and shell bins of drought responsive genes in the natural soil dataset. (**C**) Shoot iron content of Col-0 plants grown under watered and drought conditions in CHAP soil at ambient pH (5.5) and in CHAP soil amended with lime to raise the pH to 8.0 (*n* = 3). (**D**) Shoot iron content of various plant species grown under watered and drought conditions in soil (*n* = 16-20). (**E**) Total chlorophyll content in Col-0 seedlings grown on Fe limiting MS and LWA media at different strengths, 4X (LWA), 3X, and 2X (*n* = 6). (**F**) Total chlorophyll content of Col-0 seedlings grown under +Fe and -Fe conditions on different media types (*n* = 6). LWA (1X agar) is the 4X LWA recipe with 1X agar and LWA (1X MS) is the identical media formulation as MS except with 4% agar. The final two boxplots are MS and LWA with FeEDTA substituted with FeCl_3_ at +Fe concentration. (**G**) Total chlorophyll content of Col-0 and *drip1,drip2* seedlings grown on MS-Fe and LWA-Fe media (*n* = 3, error bars represent the standard error). Data in D,E,F,G were analyzed using ANOVA followed by Tukey’s test where letters indicate statistically significant differences between conditions. Data in A,C,H were analyzed with two-tail t-tests, statistical significances indicated by ∗∗∗ p < 0.001, ∗∗ p < 0.01, ∗ p < 0.05, and ns, not significant.

**Figure S5.**
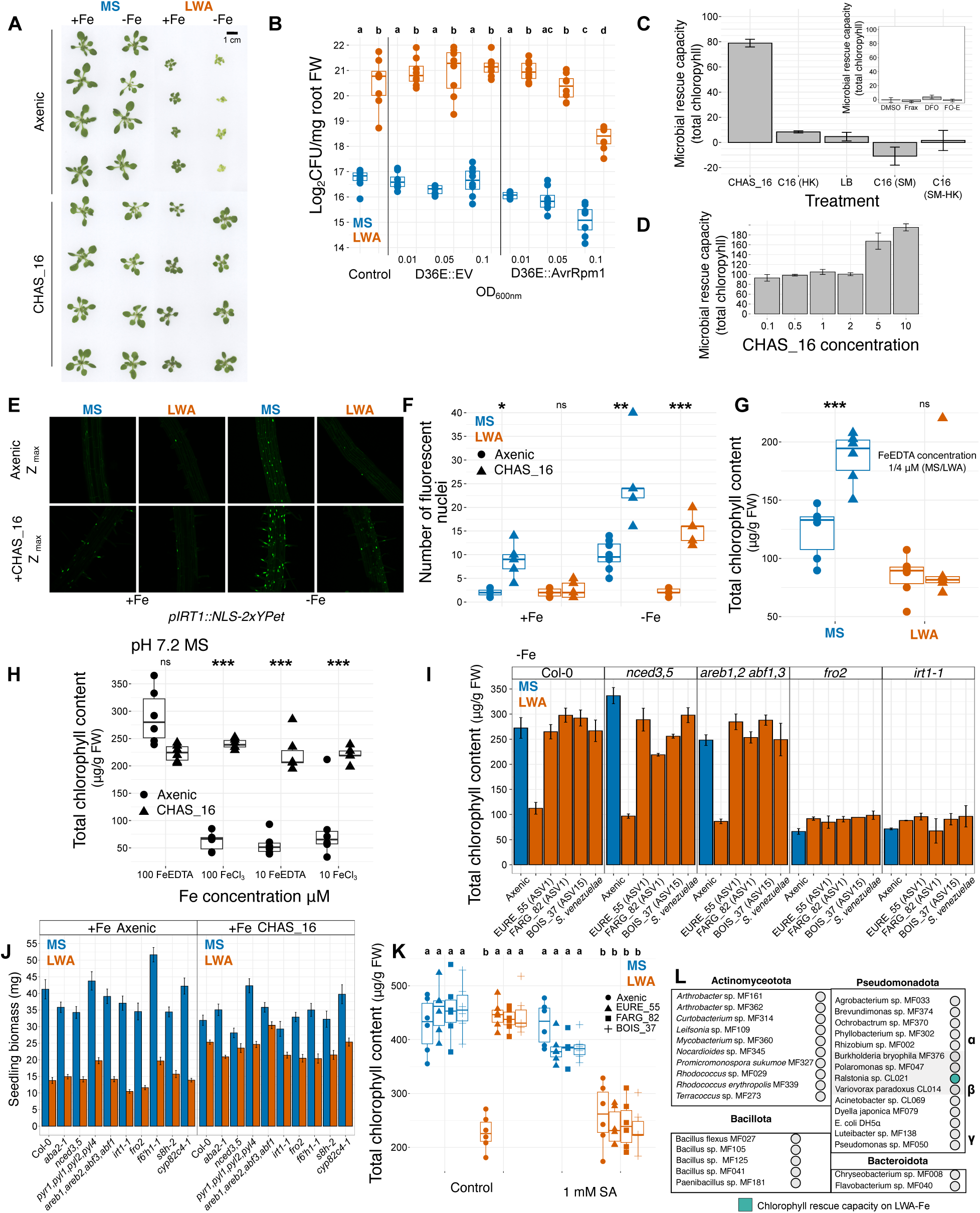
Streptomyces benefit requires a live bacterium and augments the iron starvation response, Related to Figure 5. (**A**) Representative images of Col-0 plants grown on control (MS) or drought-mimicking media (LWA) at replete (+Fe) and limiting (-Fe) iron concentrations. Plants were grown under axenic conditions or in the presence of CHAS_16, an exemplary ASV1 strain with biomass and chlorophyll rescue capacity. (**B**) CFU counts of CHAS_16 from Col-0 roots grown across listed conditions in the presence of a non-virulent Pto DC3000 mutant lacking all 36 effectors (D36E::EV) or D36E::AvrRpm1, in which the ETI-eliciting effector AvrRpm1 is restored (*n* = 8). (**C**) Chlorophyll rescue capacity of CHAS_16 requires live bacteria and is lost after heat killing and is not present in spent media (*n* = 3-6, error bars represent standard error). The inset shows that the plant coumarin fraxetin and the *Streptomyces* derived siderophores desferrioxamine and ferrioxamine E, are not sufficient to rescue chlorophyll production of Col-0 plants on LWA-Fe media (*n* = 6). (**D**) Chlorophyll rescue capacity of CHAS_16 is stable across a range of concentrations and can be enhanced at higher doses (*n* = 3-6, error bars represent standard error). A concentration of one is equal to 40 mg of mycelial mass suspended in 1 mL of MgCl_2_. (**E**) IRT1 promoter activity in roots and (**F**) the number of fluorescent nuclei of the pIRT1 reporter line grown on MS or LWA media at replete and limiting iron concentration with and without CHAS_16 (*n* = 6-8). (**G**) Total chlorophyll content of Col-0 seedlings grown on MS and LWA media at very limiting iron concentrations with and without CHAS_16 (*n* = 5-6). (**H**) Total chlorophyll content of Col-0 seedlings grown on high pH media with different sources of Fe and at different concentrations with and without CHAS_16 (*n* = 5-6). (**I**) Four additional *Streptomyces* with high chlorophyll rescue capacity of Col-0 on LWA-Fe maintain this capacity on mutants abrogated in the ABA pathway but fail to rescue *irt1* and *fro2* mutants (*n* = 6, error bars represent the standard error). (**J**) The biomass rescue capacity of CHAS_16 under LWA+Fe conditions is maintained across all tested mutants (*n* = 6, error bars represent the standard error). (**K**) The chlorophyll rescue provisioned by three additional *Streptomyces* with high chlorophyll rescue capacity of Col-0 on LWA-Fe is abolished when plant immunity is artificially activated (*n* = 5-6). (**L**) Out of 30 tested non-*Streptomyces* bacteria only one rescues chlorophyll production of Col-0 plants on LWA-Fe (*n* = 3-6). Statistical significances indicated by ∗∗∗ p < 0.001, ∗∗ p < 0.01, ∗ p < 0.05.

**Figure S6.**
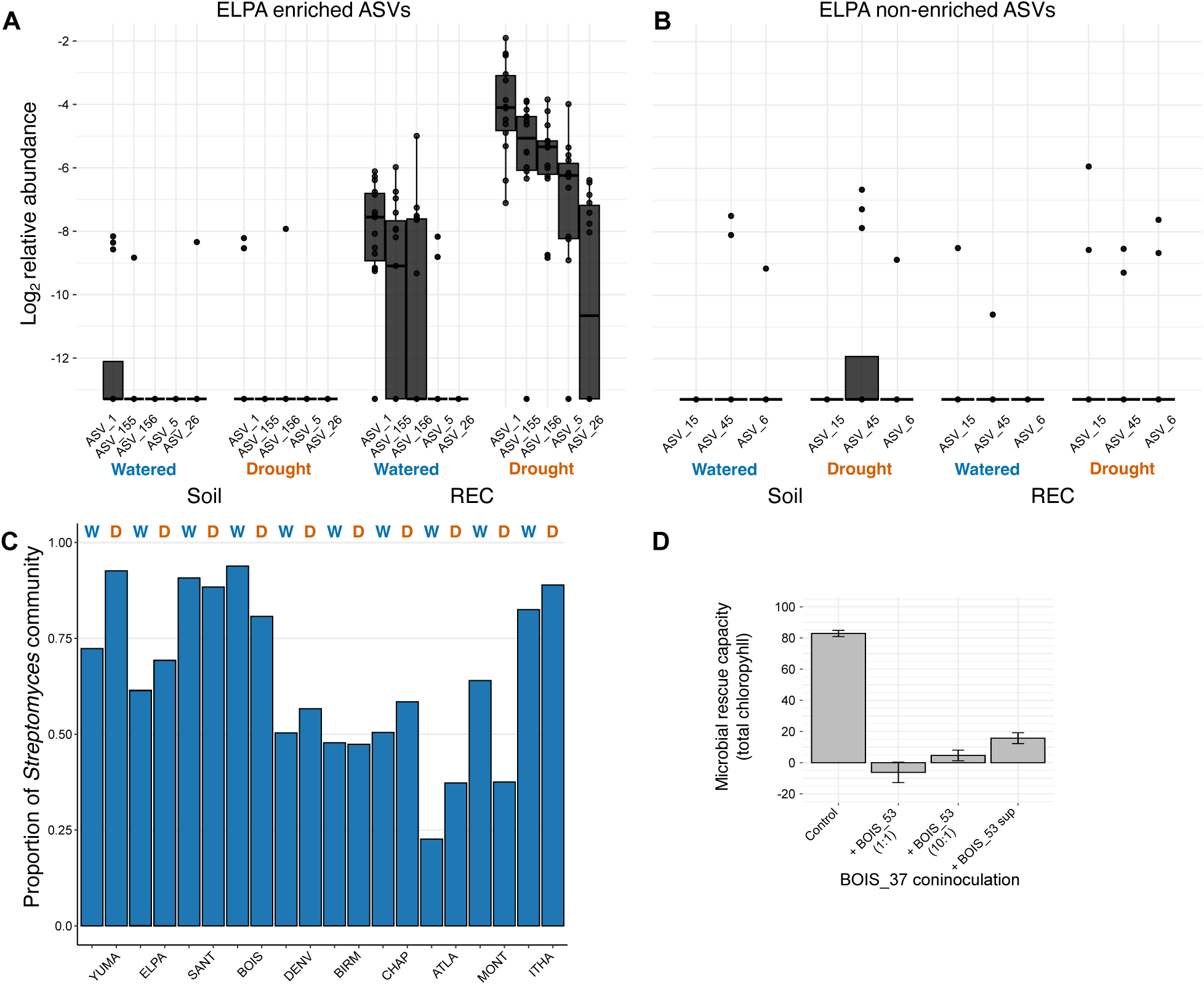
The generation of *Streptomyces* SynComs, Related to Figure 6. (**A**) The abundance profile of *Streptomyces* ASVs identified as drought enriched in the REC at ELPA (*n* = 12-15). (**B**) The abundance profile of *Streptomyces* ASVs identified as drought non-enriched in the REC at ELPA (*n* = 12-15). Isolates exactly matching the ASV sequences of E and NE ASVs were used to generate SynComs for each of the 10 selected soils. (**C**) The proportion of the total *Streptomyces* community found in the REC under watered and drought conditions represented by the ASVs used to generate SynComs. (**D**) BOIS_53 inhbits the chlorophyll rescue capacity of BOIS_15 when inoculated at ten-fold lower abundance and by way of supernatant derived from BOIS_53 (*n* = 3, error bars represent the standard error). Significance in E,G,I determined by Wald tests with FDR adjusted p values. Statistical significances indicated by ∗∗∗ p < 0.001, ∗∗ p < 0.01, ∗ p < 0.05.

## Methods

### Experimental model and study participant details

The plants used included *Arabidopsis thaliana* (Arabidopsis)*, Solanum lycopersicum* (tomato, micro-tom)*, Oryza sativa* (rice, Nipponbare), *Triticum aestivium* (CWRS), *Zea mays* (Sugar Buns F1), *Brassica napus* (Westar), and *Pisum sativum* (CDC Amaraillo). The Arabidopsis genotypes used included Col-0 (wild-type) and the following mutants, all in the Col-0 background: nced3, *nced3 nced5, pyr1 pyl1 pyl2 pyl4, aba2-1, areb1 areb2 abf3 abf1, drip1 drip2, bts-3*, *btsl1 btsl2, IMA1ox, pIRT1::NLS-2xYpet*.

Plants were grown in two different settings: in pots filled with natural soil or peat-based substrate in a growth chamber; in square 12 × 12-cm plates on agar-based media (see Agar-based plant experiments below). In both settings, seeds of all plant species were first surface sterilized (after removal of seed hulls as necessary) by agitation in 70% ethanol for 1 min, washed with sterile, distilled H_2_O (hereafter dH_2_O), agitated in 40% household bleach for 5 min, and washed 5 times with dH_2_O. Rice and tomato seedlings were placed on ½ MS media (PhytoTech Labs M404) at pH 5.7 supplemented with 0.5% sucrose and Gamborg vitamins (hereafter germination media) and incubated at 28° C in the dark for 5 days before transfer to experimental media. After sterilization, Arabidopsis, *B. distachyon, B. napus,* and *P. sativum* seeds were kept in dH_2_O at 4° C in darkness for 2 days before sowing on germination media or in pots. After growth for 7 days on germination media, Arabidopsis seedlings were then transferred to experimental media.

For agar-based experiments, Arabidopsis was grown at 21° C day and 19° C night temperatures with a photoperiod of 16 hours and 150 µmol/m^2^/s light intensity. Tomato and rice were grown at 28° C day and 23° C night temperatures with a photoperiod of 12 hours and 1000 µmol/m^2^/s light intensity. The duration of all agar-based experiments was 12 days.

For pot-based experiments, we grew plants in Cone-Tainers (Stuewe and Sons SC10U) filled with natural soil or peat-based substrate in a growth chamber at 21° C day and 19° C night temperatures with a photoperiod of 8 hours and 150 µmol/m^2^/s light intensity. The peat-based substrate consisted of 80% bark chip/peat mix, 10% sand, and 10% perlite (volume percentages). To impose a standardized drought treatment across soils with different physical properties we performed a water retention analysis on all substrates to determine the gravimetric soil moisture content after applying -0.1 (field capacity) and - 15 bars (permanent wilting point) of pressure. We measured the gravimetric soil moisture content of the starting substrate, weighed the substrate filled in each pot, estimated the dry substrate weight in each pot, and based on this value and the water retention analysis we calculated target pot weights to achieve our desired soil moisture states of field capacity and permanent wilting point. Seeds were sown on saturated substrate and covered with a clear, plastic dome to maintain humidity for 1 week. After which, the substrate was maintained at a soil moisture corresponding to field capacity for an additional week. After two weeks of benign growth conditions, experimental treatments began where, based on gravimetric soil moisture estimates, watered pots were maintained at field capacity and drought pots at permanent wilting point for 6 weeks. Pots were weighed 3 times a week and watered with H_2_O from above to maintain target weights for a duration of 6 weeks.

### Method details

#### Site selection and soil collection and characterization

We selected sites for soil collection to span the climate breadth observed in the contiguous USA. First, we downloaded bioclimatic (bioclim) data from the WorldClim dataset^112,113^, subset to include 28,338 points locations corresponding to villages, towns, and cities. We then selected locations that spanned bioclim variables 1 and 12-19 as we were most interested in capturing precipitation related variation. From these we contacted local park or conservation authorities and botanical gardens to arrange for the collection and shipment of soil (5-20 cm in depth) from locations free of any amendments or management. In total, 18 locations were selected for soil collection (Table S1). All soil was received within a span of 2 months and was held at room temperature in plastic buckets. To facilitate homogenization and remove large debris, we sieved all soils at 2 mm. Effort was made to reduce cross contamination between soils during handling by changing gloves between soils, surface sterilizing or autoclaving equipment used to handle the soils. After soil homogenization we sent aliquots to the Environmental and Agricultural Testing Service Laboratory (EATS), Department of Crop and Soil Sciences, at North Carolina State University for water retention analysis and quantification of nutrient standing stocks including C, N, NO_3_, NH_4_, P, K, Ca, Mg, S, B, Na, Cu, Fe, Al, Mn, and Zn. To quantify plant available nutrients in addition to standing stocks we used plant root simulator probes (Western Ag) incubated in each soil under watered and drought conditions for the duration of the experiment (8 weeks). The probes are ion exchange resin membranes and allow for the measurement of bioavailable NO_3_, NH_4,_ P, K, S, Ca, Mg, Al, Fe, Mn, Cu, Zn, B, Al, Pb, and Cd.

#### Bacterial community sampling

After 8 weeks of growth plants and soil were harvested for bacterial community profiling. Replicate plants in each soil X treatment combination were separated into different fractions. First, two mature and fully expanded leaves were harvested for the leaf microbiome. Next, loose soil was shaken from the entire root system followed by vortexing in dH_2_O at max speed for 10 s. Note that for very large plants sampling the entire root system was not feasible, in this case we subsampled to include first, second, and third order roots thus matching the span of root age found in the entire root system. The soil remaining in the tube after this first vortex was designated as the rhizosphere. Next, the roots were vortexed two additional times (separate tubes of dH_2_O) before being placed in new tubes with dH_2_O and sonicated (Diagenode BioRuptor, low setting 5 cycles of 30 s on 30s off). The material stripped from the root surface during sonication was designated as the rhizoplane. After sonication, roots were rinsed with dH_2_O and placed in a new tube with 3 glass beads (4 mm), these samples were designated as root endophytic compartment. Rhizosphere and rhizoplane samples were pelleted by centrifuging tubes at 6800 rpm for 5 min. From replicate unplanted pots we collected bulk soil samples. All samples were lyophilized for 48 hours prior to DNA extraction.

#### DNA extraction, PCR amplification, and sequencing

We isolated total DNA from each sample using the Qiagen DNeasy PowerSoil Pro Kit with the following modifications. Leaf and REC samples were homogenized (MP Bio FastPrep 24, 45 s) prior to input into the kit. Lysis buffer was added to individual sample tubes, mixed, and then the entire contents transferred back to the bead tubes from the kit. After DNA isolation we amplified the V3V4 region of the 16S rRNA gene using a two-step PCR protocol using dual indexed primers. In the first step, the 15 µl reaction included 3.6 µl of 5X HiFi buffer, 0.54 µl dNTPs (10 mM each), 0.9 µl DMSO, 0.36 µl KAPA HiFi Hotstart DNA polymerase, 0.9 µl each of forward and reverse primers (10 µM each), 2.7 µl mix of chloroplast and mitochondrial peptide nucleic acid (10 µM each), and 5.1 µl dH_2_O. The reaction conditions were 95° C for 5 min followed by 30 cycles of 98° C for 20 s, 75° C for 15 s, 55° C for 15 s, and 72° C for 1 min. Diluted reactions from this first step (100 fold) were used as template for the second step. In the second step, the 10 µl reaction included 2 µl of 5X HiFi buffer, 0.3 µl dNTPs (10 mM each), 0.5 µl DMSO, 0.2 µl KAPA HiFi Hotstart DNA polymerase, 1 µl each of forward and reverse primers (10 µM each), and 0.15 µl mix of chloroplast and mitochondrial peptide nucleic acid (100 µM each). The reaction conditions for the second step were 95° C for 5 min followed by 10 cycles of 98° C for 20 s, 75° C for 15 s, 55° C for 15 s, and 72° C for 1 min. The PCR products from this second step were visualized on a 1.2% (w/v) agarose gel, pooled based on band intensity, and purified 1:1 with AmPure magnetic beads (Beckman Coulter). The pooled and purified libraries were then sequenced on the Illumina MiSeq using the V3 600 cycle kit (PE, 2x 300bp) loaded at 8 pM and supplemented with 10% PhiX DNA.

#### Analysis of 16S rRNA sequencing data

After demultiplexing, we processed sequencing reads using the R package dada2^114^ version 1.24. We removed primer sequences and trimmed the end of each paired end sequence as needed to remove low quality sequences. We removed any sequences with ambiguous nucleotide assignment, with any instance of a Q-score less than 2, or with greater than 2 expected errors. We parameterized an error model and inferred sequence variants for each read prior to merging the denoised sequences and generating a sequence table. The above was performed for every individual MiSeq run to account for differences across each sequencing event. We merged sequence tables from all MiSeq runs, removed chimeric sequences and assigned taxonomy to individual ASVs using the RDP naïve Bayesian classifier (implemented in dada2) and the RDP training set 18^115^. Next, we used the R package phyloseq^116^ version 1.40 to further process our samples. We removed ASVs that were unassigned to the Bacterial kingdom, unassigned to a bacterial phylum or assigned to plastid and mitochondrial lineages. Finally, after the above filtering, we removed samples that did not have at least 500 individual sequences. To further eliminate spurious ASVs we eliminated any ASV not found in at least two samples with two reads, which retained 99.8% of the reads in the entire dataset. We then calculated alpha diversity indices in every sample (richness, Shannon’s diversity, Inverse Simpson’s diversity).

To facilitate the comparison of community composition and differential abundance testing of bacterial taxa we first simplified our dataset to include only the common ASVs. We applied a prevalence and abundance threshold where for a given subset of samples (e.g. REC at CHAP soil), ASVs had to be found in at least 20% of samples at an abundance of at least 5 sequences per sample. Differential abundance analysis was performed using the DESeq2^117^ (version 1.36) function DESeq, with the significance of differential abundance evaluated by Wald tests. P-values were adjusted for multiple testing using the Benjamini-Hochberg method to control the false discovery rate (FDR).

#### 16S copy number estimation

We used a molecular counting method^42^ to estimate 16S copy number in Col-0 roots grown under watered and drought conditions in natural soil. REC samples were collected as described above and weighed before lyophilization. Before DNA isolation we added a known number of plasmid copies carrying a sham sequence of DNA which matched the average length and GC content of a V3V4 amplicon derived from a Col-0 REC sample harvested from CHAP soil. The sham sequence was flanked by DNA complementary to the V3V4 universal primers. Plasmid number was titrated based on sample weight such that between 5-20% of all amplicons from a given sample derive from the sham sequence. The number of plasmid copies added divided by the number observed yields a conversion factor, which when multiplied by the total bacterial V3V4 16S sequences provides an estimate of the 16S copy number in each sample.

#### RNA extraction and RNA-seq library preparation

We extracted RNA from Arabidopsis leaves and roots following previously published methods. Root and leaf samples were flash frozen and stored at −80 °C until processing. Frozen tissue was ground using a TissueLyzer II (Qiagen), then homogenized in a buffer containing 400 μl of Z6-buffer; 8 M guanidine HCl, 20 mM MES, 20 mM EDTA at pH 7.0. Four hundred μl phenol:chloroform:isoamylalcohol, 25:24:1 was added, and samples were vortexed and centrifuged (20,000g, 10 min, 4 °C) for phase separation. The aqueous phase was transferred to a new 1.5-ml Eppendorf tube and 0.05 volumes of 1 N acetic acid and 0.7 volumes 96% ethanol were added. The RNA was precipitated at −20 °C overnight. Following centrifugation (20,000g, 10 min, 4 °C), the pellet was washed with 200 μl sodium acetate (pH 5.2) and 70% ethanol. The RNA was dried and dissolved in 30 μl of ultrapure water and stored at −80 °C until use.

We generated stranded mRNA-seq libraries using the KAPA mRNA Hyperprep kit for Illumina platforms. We followed the manufacturer’s protocol except for including a stubby adaptor ligation step to utilize IDT UDI primers for library amplification and indexing. Final libraries were pooled in equimolar concentration and sequenced using the NovaSeq 6000 S4 200 bp PE reagent kit.

#### RNA-seq analysis

Raw sequencing data in FASTQ format were initially assessed for quality using FastQC version 0.12.1. Adapter sequences, low-quality reads, and bases with a Phred score < 20 were trimmed using Trimmomatic^118^ version 0.36. The cleaned reads were then aligned to the Arabidopsis TAIR10 genome using BBMap version 39.13 with default settings. Alignment quality was evaluated and sorted using SAMTools^119^ version 1.21 Gene expression quantification was performed using featureCounts from SubRead version 2.0.6, with gene models obtained from TAIR10. The count data were normalized for library size and composition biases using the DESeq2 package version 1.44.0 in R version 4.3.0. DESeq2 was used to identify differentially expressed genes (DEGs) between experimental conditions. Preliminary data exploration, including principal component analysis (PCA) and hierarchical clustering, was conducted to assess sample quality and group separation. Differential expression analysis was performed using the DESeq2 function DESeq(), with the significance of gene expression differences evaluated by Wald tests. P-values were adjusted for multiple testing using the Benjamini-Hochberg method to control the false discovery rate (FDR). Genes with an FDR < 0.05 and log fold change > 1.5 were considered significantly differentially expressed. Visualization of DEGs was performed using heatmaps, volcano plots, and MA plots generated with the PHeatmap version 1.0.12 and ggplot2 version 3.5.1 packages in R. Gene ontology (GO) enrichment analysis of DEGs was performed using the clusterProfiler^120^ package version 4.12.6 and the function enrichGO to identify overrepresented biological processes, molecular functions, and cellular components. GO terms with adjusted p-values < 0.05 were considered statistically significant.

To identify gene modules associated with different conditions or traits, a Weighted Gene Co-expression Network Analysis (WGCNA) was performed on the preprocessed VST transformed gene expression data. The WGCNA procedure was carried out using the WGCNA^51^ R package (v1.69). First, a correlation matrix was computed for all genes across all samples, using the pairwise Pearson correlation coefficient to assess the relationship between gene expression profiles. A soft thresholding power (β) of 6 was selected to ensure the network adhered to a scale-free topology. The appropriate β value was determined by assessing the fit of the network to a power law distribution using the function *pickSoftThreshold* in the WGCNA package, with a target of 0.8 for the scale-free topology criterion. The adjacency matrix was then calculated by raising the absolute value of the Pearson correlation coefficients to the power of β. To construct the network, the adjacency matrix was transformed into a topological overlap matrix (TOM), which reflects the interconnectedness of genes and reduces noise from spurious correlations. A dissimilarity matrix was derived from the TOM and subjected to hierarchical clustering using the *hclust* function, with the resulting dendrogram cut at a height corresponding to a minimum module size of 20 genes.

The genes were clustered into modules based on their co-expression patterns, and module colors were assigned to each gene based on the cluster it belonged to. The number of modules was determined using dynamic tree cutting with a minimum module size threshold of 20 genes and a cut height of 9.4 which was found to balance module detection sensitivity and specificity. The modules were then characterized by their gene significance (GS) and module eigengene (ME). Module eigengenes represent the first principal component of the gene expression profiles within each module, summarizing the overall expression pattern. The module-trait relationships were assessed by correlating the module eigengenes with soil physical and chemical data to identify biologically relevant modules. The significance of these correlations was determined using the Pearson correlation coefficient and p-values corrected for multiple testing using the Benjamini-Hochberg method.

We generated literature derived sets of SA- and Fe-responsive genes. SA responsive genes were defined as those exhibiting differential expression upon foliar application with BTH from Huot et al.^121^. Fe-responsive genes were defined as the intersect of those found to be differentially expressed in three separate studies investigating the effects of iron limitation on root gene expression in Col-0^81,82,122^.

#### Wild soil BTH application and lime and ferrozine amendment

We therefore performed experiments with CHAP soil, which exhibited intermediate *Streptomyces* enrichment (Figure 3A) and thus allowed for observation of experimentally induced changes in enrichment. To maintain plant immunity during drought we foliarly applied 50 µM BTH amended with 0.02% Silwet L77 every two weeks over the course of an 8 week drought experiment in CHAP soil for a total of 4 applications. Control plants were treated with a 0.02% Silwet L77 solution at the same rate as the BTH plants. To artificially raise CHAP soil pH we added lime at an intermediate (I) and high (H) rate: CaCO_3_ (per kg soil, I=2g; H=4g) and NaHCO_3_ (per kg soil, I=1.5g; H=3g). Lime was added to soil and thoroughly mixed with soil prior to sowing seeds. To restrict iron access we added ferrozine at the rate of 10 ml of 500 µM solution to each 100ml pot filled with 95 g of moist soil.

#### qRT-PCR

Total RNA were extracted from frozen tissue using RNeasy 96 (Qiagen, 74181). On-column DNase I was used to remove genomic DNA contamination (Qiagen, 79254). Total RNAs were reverse transcribed (Superscript III, ThermoFisher 18080044) using 18bp long oligo dT (Eurofins). cDNAs were diluted tenfold and used for amplification with Power SYBR Green master mix (ThermoFisher, 4367659) and gene specific primers.

#### Fraxetin antibacterial activity

The antimicrobial activity of fraxetin (Sigma PHL89549) against single bacterial strains was assayed in liquid culture in LB. Fraxetin stocks were prepared in sterile DMSO and stored at -80° C. To reduce clumping, *Streptomyces* cultures were baffled using 10-15 4mm sterile glass beads and shaken in liquid LB media for 12-36 hours at 250 rpm at 28°C. Cultures were normalized to OD_600_ of 0.01in a final volume of 100 µL in a clear flat-bottom 96-well plate supplemented with fraxetin for a final 50 µM concentration, or DMSO negative control. The plates were then incubated at 250 rpm at 28 °C for 20 h, after which the total culture volume was combined with 100 μl BacTiter-Glo reagent (Promega BacTiter-Glo Microbial Cell Viability Assay) in an opaque plate and incubated at room temperature for 7 min. The luminescent signal was measured in a BioTek Synergy H1 microplate reader. Blank media supplemented with fraxetin or DMSO were also used to measure background luminescence and correct RLU values. Inhibition was calculated as the proportional reduction of RLU in fraxetin supplemented versus DMSO supplemented cultures.

#### Agar-based plant experiments

Our solid media included MS and Gamborg vitamins with pH adjusted with MES. The MS recipe for control conditions consisted of: 1% agar, 2.5 mM MES, macronutrients (10.31mM NH_4_NO_3_, 9.40mM KNO_3_, 1.13mM CaCl_2_ 2H₂O, 0.37mM MgSO_4_ 7H₂O, 0.62 mM KH_2_PO_4_), micronutrients (50 µM H_3_BO_3_, 50 µM MnSO_4_ H₂O, 15 µM ZnSO_4_ 7H₂O, 0.05 µM CuSO_4_ 5H₂O, 0.5 µM Na_2_MoO_4_ 2H₂O, 0.05 µM CoCl_2_ 6H₂O, 2.5 µM KI), Gamborg vitamins (277.5 µM myo-Inositol, 4 µM Nicotinic Acid, 2.43 µM Pyridoxine HCl, 14.82 µM Thiamine HCl). Iron was then added at 100 µM (+Fe) or 10 µM (-Fe) FeEDTA, or as noted in figures and figure legends. The LWA consisted of the above recipe multiplied 4X or as noted in figure legends. The pH of both media was adjusted to 5.7 using NaOH. To test whether different forms of osmotic stress resulted in the suppression of iron uptake, the MS was followed above and NaCl was added to a final concentration of 100 mM. Additionally, we implemented a PEG-infusion method^123^ to impose a water potential of -1.2 MPa.

For inoculation, bacterial isolates were shaken in liquid LB media for 12-36 hours at 250 rpm at 28° C. To reduce clumping, *Streptomyces* cultures were baffled using 10-15 4mm sterile glass beads. Bacterial cells were pelleted at 6,800 rpm for 10 mins, washed twice with 10 mM MgCl_2_ and normalized prior to inoculation. *Streptomyces* cultures were normalized by resuspending wet mycelial mass in 10 mM MgCl_2_ at 40 mg/ml. For all other bacteria we measured the optical density at 600 nm (OD600) and normalized to OD_600_ = 0.2. From the normalized cultures, 100 µl was spread onto solid experimental media plates prior to seedling transfer. For inoculations with multiple *Streptomyces* strains, equal volumes of normalized cultures were combined and 100 µl was spread onto solid experimental media. To test whether BOIS_53 sup alone could eliminate the rescue capacity of BOIS_37, 10 µl of sup was added to 90 µl of live BOIS_37 culture generated as described above and spread on experimental media. Solutions of iron chelating compounds were spread on experimental media prior to seedling transfer at the following concentrations (fraxetin 50 µM, deferoxamine mesylate 16 µM [Sigma D9533], ferrioxamine E 16 µM [Sigma 38266]).

To activate plant immunity prior to transfer to experimental media we treated plants with chemical elicitors or live bacteria. For chemical elicitors, 7-day old germination plates were flooded with 20 ml of filter sterilized SA or BTH solutions. Flooded plates were left at room temperature in a lit but off laminar flow hood for 5 hours and then transferred to experimental media. Control plates were flooded with sterile water and treated identically.

For live bacteria treatment, overnight LB cultures of D36E::EV or D36E::AvrRpm1 were washed twice with 10 mM MgCl_2_ and diluted to various concentrations. After transfer of seedlings to experimental media, 10 µL of bacterial culture was spot inoculated onto shoot apical meristems of each seedling. Plates were left in a horizontal position for 2 hours to promote bacterial colonization.

#### Characterizing iron starvation responses

Total chlorophyll content measurement was determined using spectrophotometry. Two shoots were harvested and pooled as one sample up to 50 mg, any liquid was removed from the sample before the FW was measured. Samples were incubated overnight in the dark at room temperature in 1ml of 80% acetone. Total chlorophyll (a + b) was calculated as follows:

Total chlorophyll (µg/g) = [20.2 (A645) + 8.02 (A663)] × extract volume (ml))/FW (g)

The iron composition of leaf tissue was determined by inductively coupled plasma mass spectrometry (ICP-MS) from 10-50 mg dried shoot tissue. Samples were first digested with 1ml H_2_NO3 (70%) and incubated at 70°C overnight. The next day samples were incubated with 0.5 ml of H_2_O_2_ (30%) additional 2h after which the samples were diluted with dH_2_O prior to analysis on an NexION 300D (PerkinElmer, Inc., Waltham, MA)

To visualize rhizosphere acidification, plants were grown on experimental media as described above (7 days on germination media, 12 days on experimental media), after which seedlings were placed onto a 1% agar plate containing 0.005% (w/v) bromocresol purple (pH 6.5 adjusted with NaOH) for 48 hr before being scanned.

To measure FCR activity, plants were grown on experimental media as described above (7 days on germination media, 12 days on experimental media). The seedlings were washed with dH_2_O five times to remove any residual media and 2 seedlings were pooled as one sample. The seedlings were incubated in the assay solution containing 0.1 mM Fe(III)-EDTA and 0.3 mM FerroZine in the dark for 1 h. FCR activity was measured by spectrophotometry (Tecan Infinite Pro 200) with the absorbance (562 nm) of the Fe(II)-FerroZine complex. The results were calculated on a root FW using the following formula: FCR activity (nmol (g h)−1) = OD562/29,800 × V (ml)/FW (g)/T (h) × 106

To measure *IRT1* promoter activity, *pIRT1::NLS-2xYpet* plants were grown on experimental media as described above (7 days on germination media, 12 days on experimental media). Seedlings were stained with a Propidium Iodide (PI) solution (200 µg/mL in Milli-Q water) for 5–10 seconds. Excess stain was removed by rinsing the seedlings with water. PI-stained images were acquired using a Zeiss LSM880 confocal microscope with a Plan-Neofluar 20x/0.50 Ph2 objective lens. Ypet fluorescence was excited using a 514 nm laser, and emission was detected through a 519–588 nm filter. PI fluorescence was excited at 561 nm, with emission collected using a 578–718 nm filter.

#### Bacterial isolation and whole genome sequencing

At the duration of the natural soil drought experiment, we collected bulk soil and REC as described above for bacterial culturing. REC samples were homogenized and serially diluted (1/10) for plating on ISP4 and water agar; bulk soil was treated similarly. *Streptomyces*-like colonies were re-streaked twice on ISP4 to obtain pure cultures. Spores were collected in 40% glycerol and stored at -80 °C.

We obtained cell lysates from each culture after 5 days of growth in LB media by incubating 10 µL of culture in 90 µL of 1:1 (v/v) dH_2_O and DMSO at 95° C for 10 min. We then performed 16S V3V4 amplification and sequencing as described above using this lysate as template and replacing the PNA with dH_2_O in each of the reactions. Approximately 85% of cultures contained a single ASV at a relative of abundance of >90% and after re-streaking to confirm pure cultures, we considered these to be clean isolates.

We first established a culture collection of *Streptomyces* isolated from Col-0 roots grown in each soil and bulk soil under drought. After sequencing the 16S V3V4 rRNA locus of each of the isolates, we then selected isolates whose sequence exactly matched those of ASVs from the microbiome dataset.

We isolated high molecular weight DNA from cells scraped off solid LB media using the Zymo Quick-DNA HMW MagBead Kit according to the manufacturer’s instructions for bacterial cells. Sample DNA was assessed by running samples on a 1% agarose gel and Qubit quantification. Isolation was repeated for samples with highly fragmented DNA or concentrations less than 20 ng/µL. DNA was resuspended in 12 µL of dH_2_O to achieve a maximum of 500 ng. Libraries were prepared using Oxford Nanopore’s Native Barcoding Kit 96 V14 (SQK-NBD114.96) and sequenced on a PromethION 2 Solo.

#### Genome assembly, annotation, and phylogeny construction

We assembled the Streptomyces genomes using Flye^124^ v2.9.2 with default parameters. Initial assemblies were visually inspected using BANDAGE^125^ to identify structural anomalies and to assess both contiguity and coverage depth. We kept genomes that exhibited either linear or circular structures and flagged those with abnormal assembly topologies either caused by low coverage or short sequence-reads. For assemblies that displayed fragmented graphs with short contigs (<10,000 bp), we applied an approach inspired by PTGAUL^126^ by filtering out short reads (<7,000 bp) prior to reassembly, which improved the continuity and quality of the final assemblies. To ensure accurate consensus polishing within Flye, we excluded assemblies with less than 30× coverage and re-sequenced these samples. Assembly completeness was assessed using BUSCO^127^ with the streptomycetales_odb10 database, which contains 1,579 conserved single-copy orthologs. Genomes with BUSCO completeness scores greater than 96% were considered high-quality. The samples with high BUSCO values (> 96%), typical genome structure (linear or circular) and high coverage (> ∼30x) were retained for downstream analyses. These high-quality assemblies were then annotated using the NCBI Prokaryotic Genome Annotation Pipeline (PGAP)^128^ with default parameters. We classified genomes using the GTDB-tk^129,130^. For phylogenetic analysis, we first obtained 1,544 single-copy BUSCO genes that were shared among at least 497 (90% of total) accessions (Table S6). Consequently, these genes were concatenated for a maximum likelihood analysis by IQ-TREE 2^131^, with each locus treated as a separate partition. A total of 1000 ultra fast bootstrap were applied. Strains identified as *Kitasatospora* sp., a sister taxon within the same bacterial family, were designated as outgroups.

#### Inhibition assays

*Streptomyces* spores were inoculated in R5 medium and grown for 48 h. The cultures were then back diluted 1:200 in 50 ml R5 medium for a total combined culture volume of 150 ml and incubated for 24-30 h until the OD_600_ reached 3. Cells were then pelleted by centrifugation at 21,000g for 30 min. The resulting supernatant was filtered with a 0.45 μm PES membrane vacuum filter and then concentrated using 100 kDa cut-off Amicon concentrators until reaching a final volume of 3 ml. The concentrated supernatant was run over an Econo-Pac 10DG desalting column (Bio-Rad), aliquoted and stored at −80 °C until use.

Target strains were shaken at 250 rpm at 28 °C in TSBY medium (1L: TSB without dextrose [Sigma T3938], 30 g; sucrose 103 g; yeast extract 10 g). Target strains were diluted in TSBY to an OD_600_ of 0.01 and 90 μl was transferred to a 96-well plate. Three wells of each target received 10 μl of a particular sup and three wells received sterile PBS buffer as a negative control. The plates were then incubated at 250 rpm at 28 °C for 20 h, after which the total culture volume was combined with 100 μl BacTiter-Glo reagent (Promega BacTiter-Glo Microbial Cell Viability Assay) in an opaque plate and incubated at room temperature for 7 min. The luminescent signal was measured in a BioTek Synergy H1 microplate reader. Blank media supplemented with sup and PBS were also used to measure background luminescence and correct RLU values. We performed two biological replicates of the protocol described above for each combination of sup producer and target. Inhibition was calculated as the proportional reduction of RLU in sup supplemented versus PBS supplemented cultures. Reductions of 90-100% were considered inhibitory and visually verified.

### Quantification and statistical analysis

All statistical analyses were performed in R using function and packages listed above. Data were analyzed using two tailed t-tests, one and two way ANOVA, PERMANOVA, and Wald tests. Lowercase letters indicate statistically significant differences between different conditions resulting from ANOVA followed by Tukey’s test. Statistical significances indicated by ∗∗∗ p < 0.001, ∗∗ p < 0.01, ∗ p < 0.05, and ns, not significant.

**Table S1. Soil attributes, related to Figure 1**.

**Table S2. Primers used, related to Figure 1**.

**Table S3. Root and leaf DRGs across localities, related to Figure 2**.

**Table S4. SA- and Fe-responsive gene sets, related to Figure 2**.

**Table S5. Media recipes, related to Figures 4, 5, and 6.**

**Table S6. BUSCO gene ID used for phylogeny, related to Figure 6**.

## Notes

### Competing Interest Statement

The authors have declared no competing interest.

### Summary of Updates

Updated supplemental figures and changes to main text.

